# Slice-PASEF: Maximising Ion Utilisation in LC-MS Proteomics

**DOI:** 10.1101/2022.10.31.514544

**Authors:** Ludwig R. Sinn, Lukasz Szyrwiel, Justus Grossmann, Kate Lau, Katharina Faisst, Di Qin, Florian Mutschler, Luke Khoury, Andrew Leduc, Markus Ralser, Fabian Coscia, Matthias Selbach, Nikolai Slavov, Nagarjuna Nagaraj, Martin Steger, Vadim Demichev

## Abstract

Quantitative mass spectrometry (MS)-based proteomics has become a streamlined technology with a wide range of usage. Many emerging applications, such as single-cell proteomics, spatial proteomics of tissue sections and the profiling of low-abundant posttranslational modifications, require the analysis of minimal sample amounts and are thus constrained by the sensitivity of the workflow. Here, we present Slice-PASEF, a mass spectrometry technology that leverages trapped ion mobility separation of ions to attain the theoretical maximum of tandem MS sensitivity. We implement Slice-PASEF using a new module in our DIA-NN software and show that Slice-PASEF uniquely enables precise quantitative proteomics of low sample amounts. We further demonstrate its utility towards a range of applications, including single cell proteomics and degrader drug screens via ubiquitinomics.

## Introduction

Data-independent acquisition (DIA) has become a cornerstone in modern LC-MS-based proteomics due to its reproducibility, high proteomic depth and excellent quantitative performance^1,2^. A key advantage of DIA is its acquisition speed, which enables high-throughput proteomics of large and biologically diverse cohorts^3–5^. Likewise, DIA has become a suitable method for the analysis of low sample amounts, with applications ranging from single-cell proteomics (SCP) and spatial tissue analysis via deep visual proteomics, to the analysis of post-translational modifications (PTMs) with low stoichiometries^6–8^. However, DIA methods used for such applications are necessarily a compromise, as higher throughput and lower sample amounts typically reduce the achievable sensitivity and quantitative precision^9–12^. A major increase in LC-MS sensitivity came with the introduction of dia-PASEF acquisition on timsTOF mass spectrometers, wherein adding an extra dimension for ion collection and separation - based on the ion mobility in a gas - resulted in enhanced ion utilisation^13^. Subsequently, several sensitivity enhancements and further developments of dia-PASEF^14–17^ have enabled a range of new applications of proteomics, including the analysis of PTMs as well as SCP and subcellular proteomics^18–21^.

In this work, we present Slice-PASEF, a DIA method that splits the precursor ion space into quasi-continuous ‘ion mobility slices’, which are then rapidly sampled through mass spectrometric analysis. This scan mode directs up to 100% of available peptide ions to fragmentation while maintaining high acquisition speeds suitable for high-throughput data acquisition. Benchmarking Slice-PASEF, we demonstrate that it boosts the proteomic depth and quantitative precision when analysing low sample amounts. We then showcase the benefits of Slice-PASEF in a range of applications, including high-throughput single-cell proteomics and the study of global protein ubiquitination upon degrader drug treatment. Slice-PASEF will be the method of choice for scenarios where maximum analytical precision and sensitivity is key.

## Results

### Slice-PASEF: optimal slicing of the precursor ion space

In PASEF (Parallel Accumulation - SErial Fragmentation) methods, ion isolation (i.e., ion trapping) and sampling is achieved in parallel using a segmented trapped ion mobility spectrometry (TIMS) system that accumulates incoming ions and subsequently releases them according to their mobility in a gas to the guiding Q1 quadrupole (Q1). The ions selected by the quadrupole are fragmented in the collision cell, with fragment ion mass-to-charge (*m/z)* values being recorded by the time-of-flight (TOF) analyser, enabling peptide identification^22^.

Correlating peptide precursor ion *m/z* values with the ion mobility, one obtains a characteristic ion map with *m/z* and ion mobility proportionally changing for ions of a given charge state *z*. Considering peptide ions with a charge of +2 or +3 and a similar ion mobility, we note that these are distributed over an interquartile range of about 70 *m/z* in the *m/z* dimension (Fig. S1). This allows tuning the Q1 to optimally guide released precursor ions with a given ion mobility from the TIMS analyser to the collision cell and detector regions. The set of Q1 isolation widths corresponding to the release of all precursors from the TIMS device (which spans a certain ion mobility range) - forming a set of *m/z* * ion mobility windows - is referred to as ‘frame’ and can feature multiple distinct Q1 isolation widths.

In the classical dia-PASEF implementation, each frame features a low number (typically two to five) of predefined, non-overlapping, and moderately narrow Q1 isolation widths (e.g., 10-50 *m/z*), with a full DIA cycle comprising a series of different frames, to cover the precursor *m/z* range of interest (Fig. 1a)^13^. We reason that the requirement for the isolation widths to not have an overlap likely stems from the limitations of early DIA software, which was originally designed to handle data without an ion mobility dimension. Together with the shape of the precursor ion space to be sampled, the selection of possible Q1 isolation ranges is therefore constrained, eventually limiting the method’s potential.

**Fig. 1:**
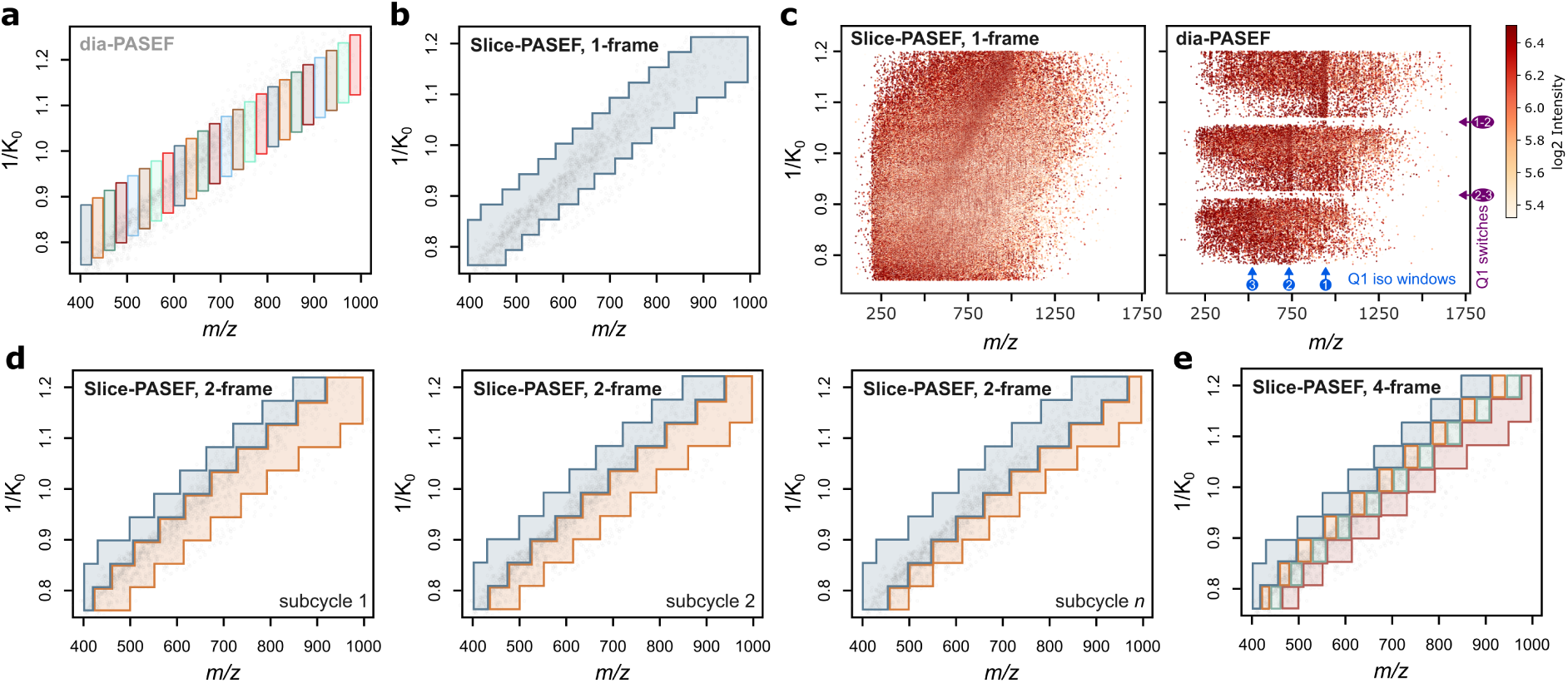
Visualisation of Slice-PASEF methods a) A common dia-PASEF method ion isolation scheme consisting of 3x8 frames of 25 *m/z* width per DIA cycle. The Q1*IMS isolation windows of each frame are highlighted with a distinct colour. b) 1-Frame Slice-PASEF method (1F) ion isolation scheme consisting of a single PASEF frame. c) Visual comparison of dia-PASEF with 1F Slice-PASEF fragment ion heatmaps from a single TIMS cycle (‘frame’), extracted by AlphaTIMS^26^. For dia-PASEF, the Q1 isolation windows (blue) as well as Q1 switches (purple) are indicated. d) 2-Frame Slice-PASEF method (2F), with shifting frame boundaries between subcycles. e) A subcycle of a 4-Frame Slice-PASEF method (4F) ion isolation scheme. Each panel indicates a full DIA cycle covering the entire *m/z*\*IM precursor ion space under investigation following an MS1 scan. Within each (sub)cycle, the precursor ion space is split into ‘diagonal’ slices, with each slice corresponding to a single PASEF frame and consisting of a number of Q1 quadruple isolation windows, which can overlap in the *m/z* dimension. Different frames are highlighted with different colours. Note that ion isolation boundaries of each frame can be shifted in the following (sub)cycles (i.e., as in the subcycles shown in panel d).

We speculated that an optimised data acquisition scheme could overcome this limitation of dia-PASEF. Specifically, if any Q1 isolation window could be used at a given ion mobility value in a frame, its width and position could be placed optimally. To explore this idea, we implemented a new module in our DIA-NN software, featuring a data representation wherein each frame is encoded as a set of signal peaks that are annotated not just with intensity, *m/z* and ion mobility, but also the corresponding Q1 isolation window boundaries (Methods). We optimised the Q1 window placement by utilising this new module, yielding a family of acquisition methods coined ‘Slice-PASEF’. In Slice-PASEF, the precursor ion space is ‘sliced’ into distinct segments that are acquired in separate frames while allowing for flexibly defining Q1 isolation windows depending on the nature of the experiment.

First, we explored how the method’s sensitivity can be maximized. We hypothesised that the resolution of the ion mobility separation coupled with recent data processing advances^14^ can essentially enable all-ion-fragmentation MS/MS DIA acquisition^23^. In what we refer to as the 1-Frame (1F) Slice-PASEF method, the whole precursor ion space except singly charged ions is sampled in quasi-continuous ion mobility slices using a single frame, wherein the Q1 is set to 50-300+ *m/z* window width with positions varying in dependence of the ion mobility (Fig. 1b). Note that with dia-PASEF all ions eluted from the TIMS device that fall outside the sampled *m/z* region at a given moment in time are lost, which constitutes a limitation in sensitivity. While this effect can partially be alleviated by tailoring the Q1 isolation widths to better capture precursor ions of interest^18^, 1F Slice-PASEF methods can be designed to lift this limitation. Interestingly, comparing raw data obtained from dia-PASEF and 1F Slice-PASEF, we further note that the latter extenuates the signal loss observed during quadrupole switches in dia-PASEF (Fig. 1c). Eventually, our method achieves a 100% MS/MS duty cycle equating to theoretically maximal MS/MS sensitivity, given the inevitable loss of analyte during ionisation and ion transmission, just as it has been shown in principle by Mann and colleagues^13,24^. Additionally, 1F Slice-PASEF allows for very short DIA cycle times. For example, at 100 ms TIMS fill time, fragmentation of all precursor ions is achieved with a single frame in 100 ms, as opposed to 800 ms (that is, 8x100 ms) for a typical dia-PASEF scheme (Fig. 1a), resulting in more data points recorded per peak and thus improved quantitation precision.

Next, we asked how we could increase the method’s specificity and decided to trade in sensitivity by dividing the precursor ion space into two segments. Further, in the resulting 2-Frame (2F) Slice-PASEF method, we leverage the overlapping isolation window concept first introduced by MacCoss and colleagues^25^ with the difference of further leveraging the TIMS in addition to the Q1 dimension. Specifically, we split the DIA cycle in a series of subcycles, wherein each subcycle features two frames with a subcycle-specific *m/z* boundary that may shift with every other subcycle (Fig. 1d). Notably, the boundaries can be optimised for splitting the precursor ion space of a given sample to enhance signal deconvolution by DIA-NN.

While 2F Slice-PASEF offers improved deconvolution of the raw data, the spectra acquired can still be relatively complex due to frequent co-isolation of co-eluting peptides which may cause highly chimeric spectra. Therefore, we explored how we can enhance selectivity in Slice-PASEF further by increasing the number of frames in a subcycle – at the cost of sensitivity due to the reduction of MS/MS duty cycle. In this work, we benchmarked the 4-Frame (4F) version of Slice-PASEF, which similarly to the 2F method utilises window boundaries that shift between subcycles. This method is designed to still provide up to twice higher MS/MS signal intensity than typical dia-PASEF schemes optimised for sensitivity^6^ (Fig. 1e). Note that Slice-PASEF frames and subcycles can be repeated several times or combined to achieve the desired balance between specificity and sensitivity, e.g., by alternating 1-Frame and 2-Frame modes, with DIA-NN then merging or deconvoluting the data originating from neighbouring frames^14^. Along with this manuscript, we also supply an application (R, Shiny) to optimally design Slice-PASEF methods customized for a given instrument and sample type based on a dia-PASEF LC-MS raw data file as input (Data availability).

In summary, in conjunction with data processing with DIA-NN (Methods), Slice-PASEF allows for optimal method design, unlocking the full analytical potential of TIMS-enabled DIA by providing precise, flexible, and efficient ion sampling strategies that can be matched to the experimental need through custom precursor isolation window and DIA cycle design.

### Slice-PASEF increases sensitivity in the proteome analysis of low sample amounts

For an initial benchmark of Slice-PASEF, we assessed its sensitivity using a dilution series of a commercial K562 cell line whole proteome tryptic digest standard and compared the results with dia-PASEF. We chose analytical flow chromatography (0.5 ml/min, 5 min gradient) coupled to a first-generation timsTOF Pro mass spectrometer. While analytical flow rate chromatography is not a preferred system for sensitive proteomics due to the high sample dilution, it is a convenient choice for conducting comparative benchmarks of acquisition methods and has the advantage of highly reproducible chromatography and high ion spray stability^3^. We compared Slice-PASEF to a typical 8-frame dia-PASEF scheme featuring 25 Da isolation windows and 12.5% MS/MS duty cycle, which we have optimised for this analytical flow rate platform^20^, and which is similar to the scheme proposed by Brunner and colleagues for single-cell dia-PASEF^6^. We observed that Slice-PASEF yields a substantial increase in peptide and protein group identification numbers specifically with low injection amounts, particularly when using the high MS/MS duty cycle schemes (Fig. 2a). For example, compared to dia-PASEF, the 1F Slice-PASEF scheme identified 84% more precursors and 46% more protein groups from 10 ng of K562 digest. The improvement in proteomic depth was achieved via the identification of low-abundant peptide ions that were missed by dia-PASEF (Fig. 2c, left). Jointly detected peptide ions were observed with a 6.2-times average signal with 1F Slice-PASEF (Fig. 2c, right). As expected for a high-sensitivity method, Slice-PASEF also attained better quantitative precision due to a higher signal-to-noise ratio and more data points-per-peak (Fig. 2b,d). For example, the 1F method was able to precisely quantify (median coefficient of variation (CV) < 10%) 3.15-times more protein groups than dia-PASEF from 10 ng acquisitions (Fig. 2a).

**Fig. 2:**
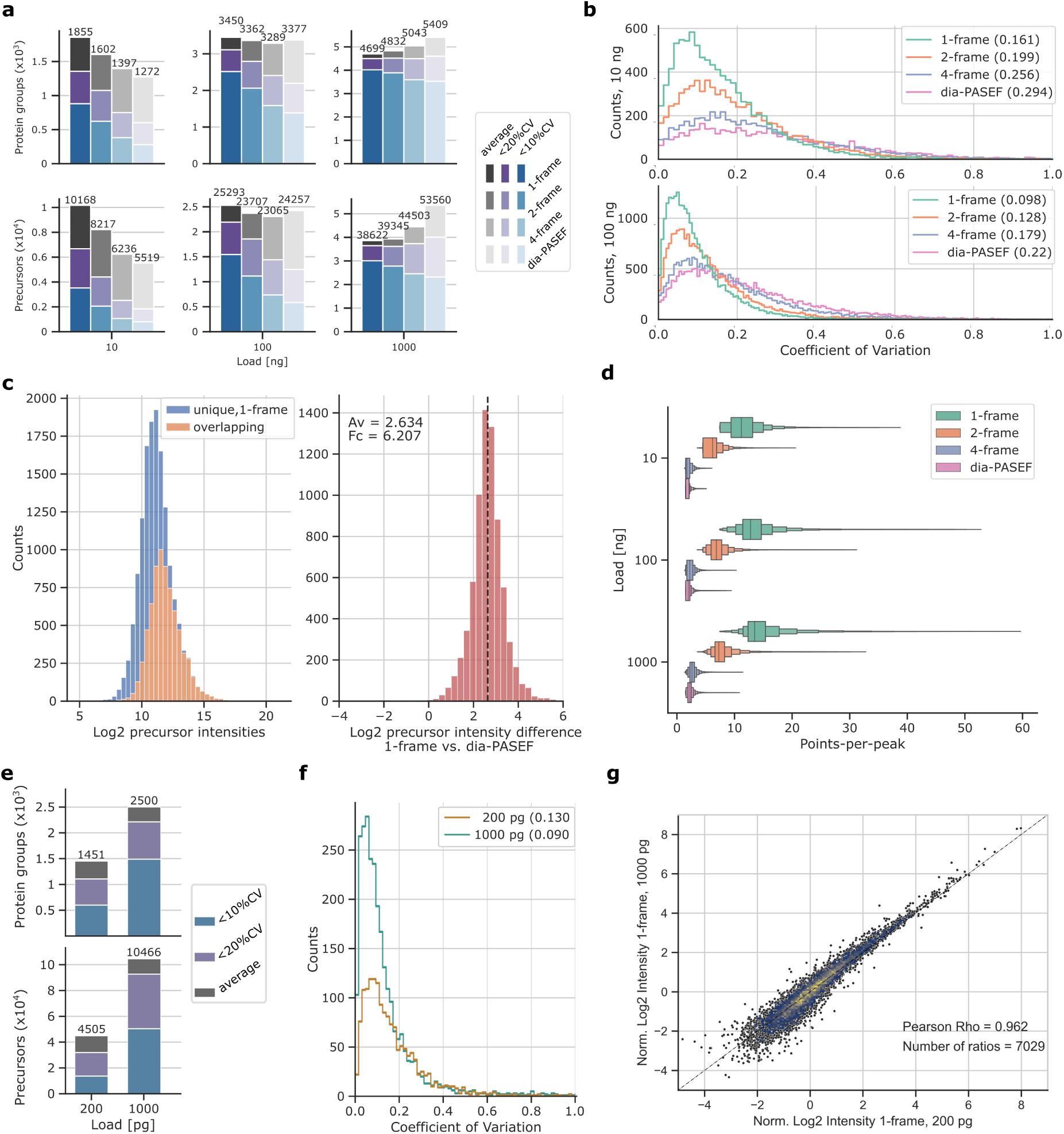
Benchmarking Slice-PASEF on analytical and fast micro-flow rate LC-MS Slice-PASEF methods compared to dia-PASEF in proteomic experiments utilising analytical flow (a-d) and fast microliter flow rate chromatography (e-g). a) Number of protein groups (upper panel) and precursors (lower panel) identified from K562 cell line tryptic digest (n=3) using different acquisition methods. Average values are indicated in grey, numbers of analytes with a coefficient of variation (CV) below 20% and 10% indicated in purple and blue, respectively (if n >= 2). b) Quantification precision, expressed as distribution of CVs for precursor quantities using 10 and 100 ng sample loads. Median values are indicated in the legend insets. c) Distribution of log_2_-transformed precursor intensities in 1F Slice-PASEF using 10 ng sample loads with identifications unique to 1F in blue, and shared with dia-PASEF highlighted in orange (left panel). Distribution of log_2_-transformed intensity ratios for precursors shared between both methods (right panel). d) Points-per-peak for each method and across three sample loads shown as Boxenplot. Median and 50% IQR are highlighted by the central box while each quantile further outwards represents half of the remaining fraction. e) Number of protein groups (upper panel) and precursors (lower panel) identified from 1 ng and 200 pg HeLa cell line tryptic digest proteome standard (n = 4) on average shown in grey, and with quantities below 20% and 10% CV in purple and blue (if n ≥ 2). f) CV distributions for protein group quantities with median values indicated in legend inset. g) Normalised log_2_-transformed intensities plotted for precursor identifications shared between 200 pg and 1 ng acquisitions. In each case, the median intensities were taken across four technical replicates.

Finally, to validate the precursor quantities obtained for ultra-low sample injection amounts given an analytical flow-rate separation (10 ng), we plotted the respective log_2_-transformed precursor quantities against those of the same precursors obtained from acquisitions using 100 ng as a reference (Fig. S2). In addition to a significantly higher number of precursors detected from 10 ng of the proteome digest standard, the 1F method showed a higher correlation between 10 ng and 100 ng sample quantities, indicating better quantitative accuracy. Notably, even the correlation between 1F Slice-PASEF * 10 ng and dia-PASEF * 100 ng, with these being from different DIA-NN searches, was higher than between 10 ng and 100 ng both acquired in dia-PASEF mode.

### Slice-PASEF facilitates high-throughput proteome analysis of single cell-level amounts

The field of single-cell-proteomics (SCP) is challenged by very low sample amounts and therefore predominantly depends on the most sensitive yet slow and delicate setups, like nano-flow liquid chromatography combined with SCP-tailored mass spectrometers such as the timsTOF SCP^6^. We reasoned that the gain in sensitivity via Slice-PASEF should allow increasing throughput for SCP applications and probed its merits using an EvoSep One system with a fast and robust 200 samples-per-day (SPD) microliter-flow rate setup connected to a timsTOF Pro2 instrument (Bruker) on a diluted HeLa cell line proteome standard digest of 0.2 and 1 ng using the most sensitive 1F Slice-PASEF method. From 0.2 ng sample, which roughly corresponds to a peptide amount of a single HeLa cell^27^, we quantified, on average, 4,505 precursors and 1,451 protein groups (Fig. 2e). The respective median CV was 13% on the protein level, from four replicate injections and only 9% when considering 1 ng load (Fig. 2f). Notably, the normalised log_2_-transformed quantities of jointly detected precursors obtained from 0.2 ng and 1 ng correlated with a Pearson correlation coefficient of 0.962, validating quantitative performance for single cell-level sample amounts (Fig. 2g).

Next, we turned to single cells isolated by fluorescence-activated cell sorting (FACS) as is routinely applied in single-cell workflows. For our test, we decided on the established U2OS Osteosarcoma cell line from which single cells were collected and processed in 384-well plates. We reasoned that due to the higher sensitivity from improved ion handling, 1F Slice-PASEF should yield a better proteome coverage, and generally higher fragment ion signals than dia-PASEF. Indeed, we quantified a median of 1,732 (+50%) proteins and 8,064 (+55%) precursors from ten individual FACS-sorted U2OS cells per method (Fig. 3a,b). Fragment ion signals increased more than 7-fold while the intact precursor signals in MS1 remained similar between both methods (Fig. 3c). Due to a stronger MS2 signal, the obtained precursor quantities displayed greater analytical precision (median CVs of 19.5% in 1F Slice-PASEF compared to 34.7% with dia-PASEF), and more low abundant precursors could be detected leading to slightly higher precursor-per-protein group counts (Fig. S3). Consequently, with Slice-PASEF we obtained deeper and more consistent coverage of a single-cell proteome while including more proteins of lower abundance, achieving an overall improved quantitative precision across all abundance quantiles (Fig. 3d-g).

**Fig. 3:**
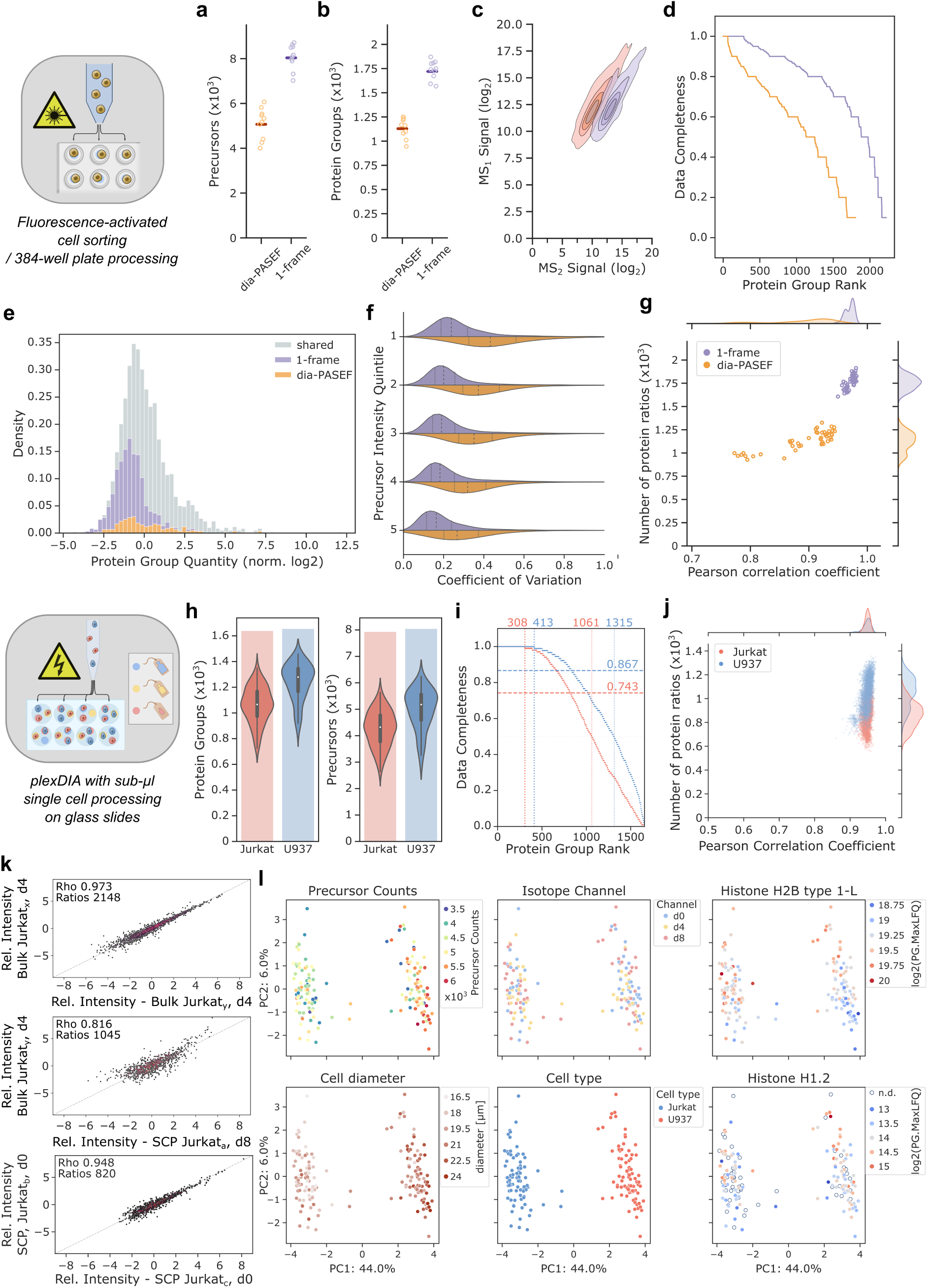
Application of 1-Frame Slice-PASEF in common single-cell proteomics workflows. a-g) Single-cell proteomics on FACS-sorted U20S cells (ten cells per method). Colour code for cell methods as in panel (g). a-b) Comparison of precursor and protein group identifications using dia-PASEF or 1-Frame (1F) Slice-PASEF. c) Ion signal intensities (log_2_-scale, Top1 precursor intensities per protein group shared between both PASEF methods) in survey (MS1) and fragment ion (MS2) scans, for both methods, shown as contour plots using five levels and scaled per method. d) Data completeness for all protein groups detected per method across the ten LC-MS runs. Protein groups were ranked by completeness per method. e) Histogram on intensity distribution of quantified protein groups. Shown are those identification unique for each method - dia-PASEF in orange and Slice-PASEF in purple - and those shared in grey. f) Quantitative precision expressed as coefficient of variation across five precursor intensity quantiles (dia-PASEF: 1502 / 1F Slice-PASEF: 1914 precursors). Dashed lines within a histogram indicate quartiles. g) Correlation of shared protein group quantities (PG.MaxLFQ) measured by Pearson correlation within replicates of each method. Histograms per axis are shown at the margins. h-l) Single-cell proteomics following the nPOP-plexDIA-approach. h) Overall protein group and precursor identifications. Violin plots show the distribution with the inner black box highlighting the IQR and the whiskers expanding to 1.5x the IQR. Medians are indicated by a white line. The bar chart in the background marks the total number of unique identifications. i) Data completeness for all protein groups detected per cell type across the measurement series, ranked from highest to least completeness. Horizontal dashed lines mark the median completeness level for each cell type dataset. Vertical dashed lines highlight 100% and 50% completeness. Numbers indicate the exact protein group counts per mark. Colour code for cell types as in panel (j). j) Relationship of shared protein group identifications with their corresponding quantitative correlation, measured by Pearson correlation, for identical cell types and per plexDIA channel. Histograms for each variable are shown at the axis margins. k) Signal correlation for protein group quantities between two bulk / a bulk and a single Jurkat cell / two single Jurkat cell plexDIA channels. Comparisons from representative channels with regards to correlation and number of protein groups are shown. l) Principal component analysis of the plexDIA dataset. Colouring is shown for variables like total precursor counts per single cell, plexDIA isotope channel, cell size, cell type, as well as quantities of two histone proteins (H2B type 1-L & H1.2).

Besides sensitivity, sample throughput is the second major challenge for SCP. To tackle this bottleneck, we turned to plexDIA^19,28^ which leverages multiple isotope channels, each encoding peptides from a different individual cell. We probed Slice-PASEF on the nPOP-plexDIA workflow using two cell lines - Jurkat and U937 - which represent lymphoid and myeloid lineages that are regularly employed in biomedical research and are characterized by rather small cell sizes, making them challenging samples for an SCP workflow^29^. The samples were run as 3-plex using mTRAQ labeling on a timsTOF SCP, with a throughput corresponding to about 135 single cells per day^19^.

For each cell type - Jurkat and U937 - and across the measurement series, we identified a total of 1,641 and 1,657 protein groups (medians 1,066 and 1279) with median data completeness of 74.3% and 86.7% for Jurkat and U937 cell proteins, respectively (Fig. 3h,i). Further, we found protein identification numbers scaling with cell size, as expected, and that most obtained protein quantities displayed low variation (median CVs of Jurkat: 24%, U937: 25%) (Fig. S4). Given that noise typically increases with lower signal, we assessed signal linearity by comparing plexDIA-channel-specific protein group quantities across the measurement series and for each cell type. First, when correlating the channel-specific quantities for Jurkat or U937 cells between replicate bulk sample measurements (Methods), we found high agreement based on Pearson correlation coefficients (PCC) of 0.973 and 0.982 from 2,148 and 2,782 protein groups (PGs), respectively (Fig. 3k). When comparing quantities between bulk and all single cell channels of matching cell types, the degree of correlation declined to a PCC_av_ of 0.810 (±0.009 s.d.) across 1018 (±129 s.d.) PGs for Jurkat cells (PCC_av_ 0.806±0.0093 s.d. across 1219±139 s.d. PGs for U937 cells). We explain this difference by a different extent of signal interferences from having a mixture of both cell types present in the bulk sample besides variance introduced during sample preparation. Notably, the signal correlation between single cell channels of identical cell type was found to be almost competitive to bulk sample measurements with a PCC_av_ of 0.949 (±0.0076 s.d.) for Jurkat and 0.948 (±0.0121 s.d.) for U937 cells (across 862 (±110 s.d.) or 1055 (±128 s.d.) PGs, respectively) (Fig. 3k, top and bottom).

Next, we asked whether we can distinguish the two cell-types by characteristic proteins. Indeed, we detected T-cell glycoprotein epsilon chain CD3E, a specific marker for the Jurkat lymphoid lineage which plays a crucial role in the adaptive immune system for T-cell activation and T-cell receptor regulation^30,31^. Further, only in U937 channels, we observed Phosphatidylinositol 3,4,5-trisphosphate 5-phosphatase 1 (INPP5D) and Tyrosine-protein kinase BTK which affect B lymphocyte development, differentiation and signaling but also link to cytokines and interleukin production thus confirming U937’s as monocyte derivative^32,33^ (Tab. S1).

Further, leveraging significant quantitative differences between the cell lines to identify cell-type-specific biological processes from a gene set enrichment analysis (GSEA), we recover processes indicative of cell lines of the immune system (Fig. S5). For instance, we extract several terms related to viral and bacterial infection, as well as endocytosis/regulation of actin cytoskeleton/phagosome enriched in U937 cells that link to cell motility and efficient phagocytosis of pathogens. Also, the GSEA returned the ribosome and protein processing in the endoplasmic reticulum, underlining that monocytic U937 cells produce immunologically relevant hormones^34,35^. For Jurkats, the recovery of mRNA surveillance pathway links to active immune-specific RNA surveillance needed for efficient triggering of viral defense, as well as the proteasome likely relating to Jurkat’s capacity to readily produce highly diverse human leukocyte antigen class I-associated peptides ^36^ (Fig. S5).

Visualising the data with principal component analysis (PCA), we also note that the quantitative proteome differences between cell types separate them into two well defined clusters, with negligible contribution from technical variables relating to sample preparation or LC-MS measurement (Fig. 3l). Instead, along the second PCA dimension, we found gradual abundance changes of histone H1.2 and H2B type 1-L which are said to roughly double along with nuclear DNA prior to mitosis thus confirming cell cycle specific proteome changes as a major driver of proteome variance within a cell (Fig. 3l)^37^. Following up, we focussed on the known cell cycle marker Nuclear ubiquitous casein and cyclin-dependent kinase substrate 1 (NUCKS1) that increases when entering S- and declines in G2/M-phases^38,39^. Since cellular growth along the cell cycle leads to relative concentration changes of certain protein subsets, e.g., a dilution of histones during G1-phase, until in S-phase, histones and nuclear DNA double prior to mitosis^40^, we analysed the distribution of NUCKS1 quantities in the context of histone protein levels as well as measured cell diameters. Here, we observed (at least) two distinct groups of U937 cells displaying a consistent abundance gradient of NUCKS1 that indeed suggests separation by cell cycle stage (Fig. S6; in line with Matzinger and colleagues^39^). Likewise, the Cyclin-dependent Kinase 1 (CDK1) which phosphorylates NUCKS1 was recovered and exhibited a correlation with NUCKS1 levels, besides other proteins with conceivable cell-cycle dependent abundance changes as suggested previously^41^ (Fig. S6).

### Slice-PASEF enables precise and sensitive Ubiquitinomics

Research focusing on post-translational modifications (PTMs) is commonly challenged by sample preparation artefacts, rapid turnover and/or chemical lability of PTMs, as well as the complexity of confident PTM identification by DIA software due to the vast search space and its combinatorial complexity^42–44^. However, the main challenge for MS-based PTM mapping lies in the low stoichiometry of many PTMs of interest. This is particularly true for protein ubiquitination that can mark proteins for proteasomal degradation. Short-lived ubiquitin-modified proteins typically display a median ubiquitination-site occupancy of less than 0.01%^45^. Yet, global mapping of protein ubiquitination is crucial for advancing both basic biological as well as drug discovery research. In the latter context, it plays a key role in validating the mode of action (MoA) of degrader drugs, such as Proteolysis targeting chimeras (PROTACS) or molecular glue degraders (MGDs)^46^. These compounds function by recruiting non-native substrates (or neosubstrates) to endogenous E3 ligases, leading to their ubiquitination and subsequent degradation by the proteasome^47^. While enrichment and detection methods for ubiquitinated peptides (ubiquitin-remnant K-GG peptides, featuring a diglycine adduct on one or more lysines) have been optimised over time, there remains a strong need to further enhance both the sensitivity and precision of quantification, particularly when working with low-input protein samples^48,49^.

To enable high-sensitivity ubiquitinomics from limited sample material, we miniaturized the wet-lab ubiquitinomics workflow and integrated it with Slice-PASEF. We enriched K-GG-remnant peptides from untreated HEK293 cells using 200 µg of protein input and diluted the eluate to simulate input amounts ranging from 50 µg down to 1 µg. We injected each sample in triplicate, alternating between 1F Slice-PASEF and dia-PASEF, starting with the lowest input. As expected, 1F Slice-PASEF outperformed dia-PASEF across all tested conditions in both K-GG peptide identifications and quantificative precision. The advantage was most pronounced at inputs of 12 µg or less, where Slice-PASEF identified up to 46% more K-GG peptides and yielded approximately three-fold more peptides with a high quantification precision (CV ≤ 20%) (Fig. 4a,b). These results highlight the clear advantage of Slice-PASEF for use in ubiquitinomics.

**Fig. 4:**
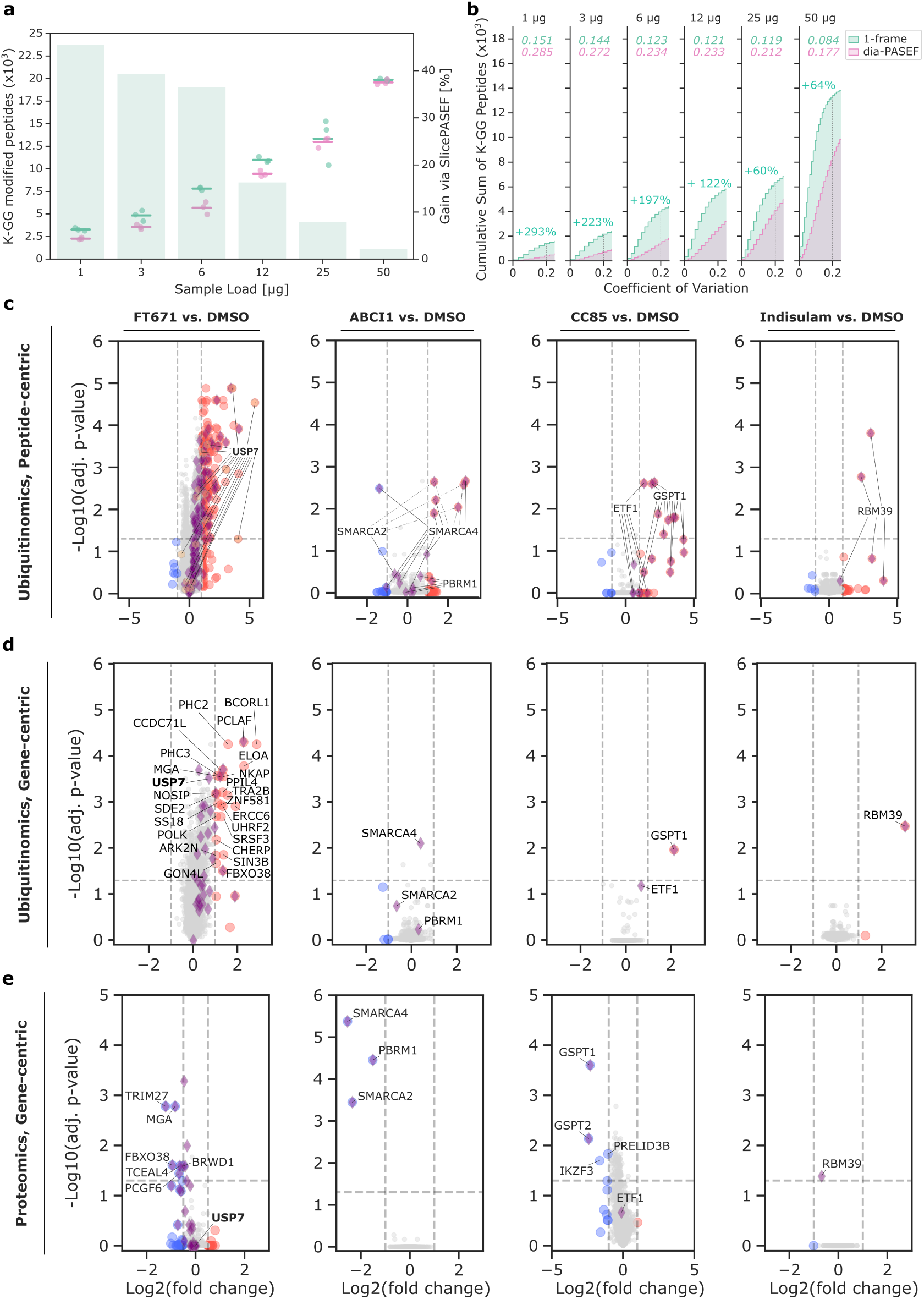
Application of Slice-PASEF for Ubiquitinomics a) Titration of proteome inputs in an antibody-based K-GG-remnant peptide profiling experiment for dia-PASEF (pink) and 1-Frame (1F) Slice-PASEF (green). The left axis displays the number of ubiquitin-remnant (K-GG) peptides. Each dot denotes the counts from one of three replicates, with a mark indicating the corresponding median. The right axis shows the percentage increase by 1F Slice-PASEF compared to dia-PASEF, indicated by the green bars in the background. b) Cumulative sum of K-GG peptides with increasing coefficient of variation (CV) for each load as in (a). Data is shown for CV values of up to 25% and only for K-GG-peptides quantified in all replicates. Median CVs of K-GG peptide quantities are indicated in the corresponding colour for both scan modes at the top of each panel. Gains in K-GG-peptide counts with CV ≤ 0.2 by Slice-PASEF are indicated in green as percentages. The vertical dashed line marks a CV of 0.2. c & d) Volcano plots showing the effect of drug treatments on the ubiquitinome on peptide-level (c), as well as gene-level (d) (Methods). Significantly up- or down- regulated gene products are marked in red or blue, respectively, and selected known targets are additionally highlighted with purple diamonds (known regulated, for FT671 according to Steger and colleagues^49^) or orange hexagons (USP7, known direct drug target of FT671) besides labelled with their corresponding gene names (in panel d). For peptide-level analysis, peptides shared between proteins are indicated with a dashed line. e) Volcano plots on the global proteome changes 6 h post treatment. Labelling keys match the other plots in panels (c) and (d).

We next profiled four ubiquitin-proteasome system (UPS)-targeting drugs using global ubiquitinomics with 1F Slice-PASEF. These included an USP7 inhibitor (FT671), a VHL-engaging PROTAC directed against SMARCA2/4 (ACBI-1), and two MGDs targeting either CRBN (CC885) or DCAF15 (Indisulam) substrate receptors^50–52^. HEK293 cells were cultured in 6-well plates and treated with each compound for 30 minutes to capture early ubiquitination events. On average, we quantified approximately 5,000 ubiquitinylated protein groups from 25,000 precursors per sample, with K-GG remnants in more than 90% of all detected precursors (Fig. S7). To connect these ubiquitination profiles with subsequent changes in protein abundance, we also quantified the global proteome after a 6-hour drug treatment using dia-PASEF with high sample amounts, leading to more than 9,000 protein groups from about 180,000 precursors (Fig. S7).

All tested drugs induced the expected effects at both the ubiquitinome and proteome levels. For instance, the USP7 inhibitor FT671 promoted the ubiquitination of USP7 itself as well as several of its reported targets, such as TRIM27^49^. We found seven additional early ubiquitinated proteins that were also downregulated in the proteome six hours after treatment, suggesting they are direct USP7 substrates. Indeed, six of those were reported before^49^ while one new hit - the Melanoma-associated antigen D4 - got residue-specific confirmation for four ubiquitination sites according to pooled public databases (Methods). Overall, our analysis is highly consistent with our previously published results, showing an overlap of about 60% (25/42) of proteins that were ubiquitinated^49^. Of the remaining 380 hits (log_2_-fold change > 0.5) that were insignificantly modulated at the protein level, 25 (6.6%) are known USP7 targets and 352 (92.6%) are predicted targets according to the UbiBrowser database. Among these, 116 (33%) are classified as high-confidence targets (p-value < 0.05)^53^.

Treatment with ACBI-1 induced the ubiquitination and subsequent downregulation of BAF complex members SMARCA2, SMARCA4 and PBRM1, confirming its expected mode of action by demonstrating intracellular ubiquitination of endogenous substrates^50^. Additionally, we validated the reported targets of both MGDs, CC885 and Indisulam (GSPT1 and RBM39, respectively), demonstrating that Slice-PASEF can facilitate the discovery of a drug’s mode-of-action and thus support studying degrader drugs in the native cellular context and in a target- and E3 ligase-agnostic manner (Fig. 4c).

## Discussion

The advent of applications that focus on hard-to-detect analytes, such as proteins from single cells or rare PTMs, has highlighted the sensitivity and throughput limitations of current workflows. In this work, we describe Slice-PASEF - a family of data-independent acquisition proteomics methods that maximize ion utilization, and thus sensitivity, by utilising trapped ion mobility separation on timsTOF mass spectrometers.

We demonstrated that Slice-PASEF outperforms dia-PASEF for proteome profiling of low sample amounts down to single-cell levels, both in terms of identification and quantification performance. Using commonly employed single-cell proteomics sample preparation techniques like FACS or dispension on glass slides as in the nPOP-plexDIA workflow, we confirm Slice-PASEF’s suitability for large-scale SCP studies given its decent proteome coverage, high data consistency, as well as increased throughput of up to about 135 single cells per day. We were able to clearly distinguish cell types based on their single-cell proteome profiles and further confirmed observations concerning cell-cycle-specific intracellular proteome variation. Meanwhile, the next generation timsTOF mass spectrometers, such as the timsTOF Ultra series, has been proven to reach even higher throughput and proteomics depth^29^, suggesting Slice-PASEF’s performance may improve in future SCP workflows, particularly when combined with cutting-edge chromatographic setups like the EvoSep One and Eno systems (EvoSep) connected to high-end separation columns (e.g. Aurora column series, IonOpticks)^54^.

Next, we illustrated the use of Slice-PASEF for studying rare PTMs, by applying it to high-sensitivity ubiquitinomics, an emerging tool for drug discovery when focusing on, i.e., PROTACs and MGDs that target proteins previously thought “undruggable”^55^. Here, our method enabled the quantification of ∼13,000 K-GG peptides with high precision (≤ 20% CV) using only 25 µg input (i.e., 2.5 - 5% of previous efforts), and notably without any drug-induced accumulation of ubiquitin-remnant peptides^49^, therefore reducing the dependency on costly ubiquitin-specific antibodies. Also, we showcased Slice-PASEF’s capacity for sensitive detection of ubiquitination events across diverse drug types such as DUB inhibitors, PROTACs, or MGDs. Here our workflow leveraged the high quantitative precision of Slice-PASEF to validate specific therapeutic profiles of the tested drugs.

Since our first release of Slice-PASEF^56^, other advanced PASEF methods were made public which also promised enhanced performance, balancing analytical specificity, sensitivity and speed of DIA measurements^15,16^. However, we note that only 1-Frame Slice-PASEF retains the maximum of attainable analyte signal at high acquisition speed, leading to comprehensive and complex data which can be handled well by our Slice-PASEF module in DIA-NN^17,57^.

We anticipate that Slice-PASEF will empower existing as well as emerging applications in proteomics that require maximal sensitivity and precision at increased sample throughput. These include spatial proteome profiling of tissues, large-scale single-cell proteomics^7,58^, and quantitative modeling approaches that benefit from enhanced precision and accuracy afforded by Slice-PASEF^59^.

We expect that future efforts will explore combining our method with scanning quadrupole acquisition, leveraging both the midia-PASEF and 2-Frame Slice-PASEF concepts. Albeit forfeiting on sensitivity compared to 1-Frame Slice-PASEF, such methods could enable high fidelity precursor identification as needed in scenarios of highly complex samples that require both high sensitivity and confidence in peptidoform identification. These include DIA-based immunopeptidomics, structural proteomics, activity-based chemoproteomics and metaproteomics^60–63^.

## Methods

### Benchmark LC-MS runs

For the benchmark analysis, a HeLa as well as a K562 cell line tryptic digest proteome standard (V6951, Promega) were diluted in 0.1% formic acid (FA) according to the manufacturer’s guidelines.

For analytical flow-rate proteomics we used an Agilent 1290 Infinity II liquid chromatography system coupled to the Bruker timsTOF Pro mass spectrometer equipped with the VIP-HESI source (3000 V of capillary voltage, 10.0 l/min of dry gas and temperature 280°C, probe gas flow 4.8 l/min and temperature 450°C). The peptide separation was performed on a Luna Omega 1.6 μm C18 100 Å 30 x 2.1 mm column heated to 60°C using a linear gradient ramping from 3% B to 36% B in 5 minutes (Buffer A: 0.1% FA; Buffer B: acetonitrile (ACN) +0.1% FA) at a flow rate of 0.5 ml/min. The column was washed by an increase to 80% B over 0.5 min at a flow rate of 0.85 ml/min, then maintained for another 0.2 min. In the next 0.1 min, the B content was reduced to 3% and 0.6 ml/min after 1.2 min and 0.5 ml/min after another 0.3 min. For the dia-PASEF method, the MS/MS precursor mass range was *m/z* 401 to 1226 and 1/𝐾_0_ 0.72 to 1.29, with 33 x 25 *m/z* windows with ramp and accumulation time 72 ms and an estimated cycle time of 0.7 s, similar to earlier work^20^. For the 1F, 2F and 4F Slice-PASEF methods the ramp and accumulation time were 100 ms and the windows setup was chosen as presented on Figure 1 (method definition files are available at https://osf.io/t2ymc/?view_only=7462fffb20e648fc83afc75d8c67e9f8). The *m/z* range was 400 to 1000 and the 1/𝐾_0_ range 0.75 to 1.2. All methods were used in the high sensitivity mode of the mass spectrometer.

For the 200 samples-per-day (SPD) method, the Evosep One system was coupled with the Bruker timsTOF Pro 2 mass spectrometer equipped with the Captive Spray source. The Endurance Column 4 cm x 150 µm ID, 1.9 µm beads (EV1107, Evosep) was connected to a Captive Spray emitter (ZDV) with 20 µm diameter (1865710, Bruker). The source parameters were kept under default settings (Capillary voltage 1400 V, Dry Gas 3.0 l/min and Dry Temp 180°C). Evotips were loaded and maintained following the manufacturer’s protocol.

Slice-PASEF and dia-PASEF methods were set up in the Bruker timsControl software (v3.0.0, analytical flow setup, and v1.1.19 68, Evosep One), by importing a text table method definition file containing the isolation window specification.

### AlphaTims evaluation on raw timsTOF data

Two timsTOF raw data files of our benchmark (see above, dia-PASEF and 1-Frame Slice-PASEF) were uploaded to AlphaTims^26^ and subsetted to fragment ion signals from a single frame (i.e., a single TIMS ion isolation and release cycle) within the LC-MS run. The filtered data was exported as csv-files for plotting (see below).

### FACS-sorted U2OS cells preparation and LC-MS measurement

Single cells were sorted into individual wells of a 384-well plate preloaded with 1 µl of sample buffer (50 mM ABC, 0.1% (w/v) n-Dodecyl β-d-maltoside (DDM), 20% (v/v) ACN) using the Mantis Liquid Dispenser (Formulatrix). Cell sorting was performed on a FACSAria III (BD Biosciences). The sample plate was sealed with aluminum foil, briefly centrifuged, and stored at -80°C. For further processing, cells were heated at 75°C for 30 min, followed by the addition of 1 µl digestion buffer (2 ng/µl Trypsin, 50 mM ABC, 20% (v/v) ACN) and an overnight digestion at 37°C. Proteome digests were analysed by LC-MS using either dia-PASEF (3×8 windows, optimised for single-cell analysis) or the 1-Frame Slice-PASEF method on a timsTOF SCP (Bruker), employing a 21-minute gradient at a flow rate of 250 nl/min on an EASY-nLC 1200 HPC with a 20 cm in-house packed column using ReproSil-Pur 120 C18-AQ 1.9 µm as solid phase (Dr. Maisch).

### Bulk proteomic sample preparation for U937 and Jurkat cell lines

U937 and Jurkat cell lines were cultured individually in RPMI medium (Sigma-Aldrich R8758) supplemented with 10% fetal bovine serum (Gibco A4766801) and 1% penicillin-streptomycin (Gibco 15140122). Cells were grown at 37°C with 5% CO_2_ and passaged approximately every two days when density reached 10^6^ cells per mL. For bulk sample preparation, cells were washed twice with 1% PBS, counted using a hemocytometer, then frozen at -80°C in a concentration of 10^6^ cells per mL in LC-MS grade water. Frozen stocks containing 100 µL were heated to 90°C in a thermal cycler for 10 minutes for lysis by mPOP^64,65^. Once cooled to 4°C, 1.3x final concentration of Benzonase Nuclease (Sigma E1014), 16 ng/µL final concentration of Trypsin Gold (Promega V5280), and 105 mM final concentration Triethylammonium bicarbonate (TEAB, pH 8.5) (Sigma-Aldrich T7408) was added. Samples were digested for 18 hours at 37°C. Digests were subsequently dried down, resuspended in 100 mM TEAB (pH 8.5), split into aliquots, and individually labeled with mTRAQ channels according to manufacturer instructions (SCIEX 4440015, 4427698, 4427700) as described by Derks et al.^19^. Final combined bulk samples contained a channel for Jurkat cells (d4), U937 cells (d8), and a third channel containing an equal mixture of the two cell lines (d0).

### Single-cell and library-bulk proteomic sample preparation for U937 and Jurkat cell lines

Cells of each cell line were washed twice with 1% PBS, counted by hemocytometer, and suspended in 1x PBS at a concentration of approximately 200-300 cells per μL. For the spectral library generation, a sample containing 100 cell equivalents of equal parts Jurkat and U937 bulk-derived peptides was labeled with mTRAQ d0. Single-cell samples using the mTRAQ 3-plex were generated using the nPOP protocol^29,66^ and the CellenONE cell sorting system (Cellenion). Cell and mTRAQ labels were permutated across the slide. Final single-cell samples were dried down in 384-well plates and stored at -80°C prior to analysis.

### plexDIA - LC-MS data acquisition of U-937 and Jurkat samples

Bulk and single-cell samples were resuspended in LC-MS grade water with 0.1% formic acid and 0.01% (w/v) n-Dodecyl β-d-maltoside (DDM). Diluted bulk samples were injected out of glass inserts in HPLC vials at approximate concentrations of 10-cell equivalents per mTRAQ channel. 3-plexDIA single-cell samples were injected out of 384-well plates. Samples for spectral library generation were run in triplicate. Peptides were separated on a 25 cm x 75 μm Aurora CSI Series nano-flow UHPLC column (IonOpticks AUR2-25075C18A) using a Vanquish Neo UHPLC (Thermo Fisher Scientific) at a flow rate of 200 nL/min. The following LC settings were selected: Direct Injection, Nano/C1/K0ap flow, maximum pressure = 1500 bar, maximum pressure change = 1000 bar, 1 μL injection volume. Loading parameters set to Fast Loading mode with Pressure Control activated and set to 1450 bar with 1.2 μL loading volume. Wash and equilibration set to Fast Equilibration with Pressure Control activated and set to 1450 bar with 4.0 equilibration factor.

The LC gradient was generated using LC-MS grade water with 0.1% FA (Fisher Scientific LS118) for mobile phase A and LC-MS grade ACN at 80% with 0.1% FA (Fisher Scientific LS122) for mobile phase B. The LC gradient began at 2.5% B, ramped to 6.5% B over 0.2 minutes, 11.5% B over 0.9 minutes, 21% B over 3.1 minutes, 31.5% B over 6.2 minutes, 40% B over 2.8 minutes, 55% B over 1.7 minutes, 95% B over 0.65 minutes, and held at 95% B for 4 minutes. Peptides actively eluted for approximately 15 minutes and total run-to-run time was approximately 32 minutes, yielding a total rate of 45 runs per day and thus up to 135 cells per day for the multiplexed samples. MS measurements were made using a timsTOF SCP (Bruker). All measurements used positive ion mode and the CaptiveSpray source (Bruker).

All dia-PASEF measurements (Jurkat and U937 library generation runs and bulk analyses) employed the following settings: 8 total PASEF frames comprising of 26 *m/z* MS2 windows with 1 *m/z* overlaps, four MS1 scans per TIMS cycle (i.e., MS1 preceding on two PASEF frames, resulting in more frequent precursor ion sampling), and accumulation and ramp times of 100 milliseconds, resulting in a duty cycle time of 1.28 seconds. The other parameters were set as follows: MS1 scan range: *m/z* 100 – 1700. MS2 scan range: *m/z* 300 – 1000. 1/K_0_ start: 0.64, 1/K_0_ end: 1.2. Collision energy settings were 20 eV at 1/K_0_: 60 and 59 eV at 1/K_0_: 1.60. Collision RF: 2000 Vpp.

Slice-PASEF SCP measurements were made using 1-Frame Slice-PASEF with slight modifications: Accumulation and ramp times were increased to 200 milliseconds, resulting in a duty cycle time of 0.41 seconds. MS2 scan range was increased to cover the lower end of the *m/z* range, specifically using the following additions: *m/z* 300 – 484 with 1/K_0_ 0.66 – 0.69, *m/z* 330 – 484 with 1/K_0_ 0.69 – 0.72, *m/z* 350 – 484 with 1/K_0_ 0.72 – 0.75, and *m/z* 350 – 484 with 1/K_0_ 0.75 – 0.78. MS1 scan range: *m/z* 100 – 1700. MS2 scan range: *m/z* 300.2 – 1000. 1/K_0_ start: 0.66, 1/K_0_ end: 1.20. Collision energy settings were 20 eV at 1/K_0_: 60 and 59 eV at 1/K_0_: 1.60. Collision RF: 2000 Vpp.

### *Ubiquitinomics - cell culture*, *drug treatments* and *cell lysis*

HEK293 cells were obtained from Cytion and cultured in Dulbecco’s Modified Eagle Medium (DMEM) (VWR), supplemented with 10% fetal calf serum (FCS) (Thermo Fisher Scientific). Indisulam, FT671 and CC885 were from MedChemExpress. ACBI-1 was kindly provided by opnMe (https://www.opnme.com). All compounds (ABCI1, CC885, FT671, Indisulam) were dissolved in dimethylsulfoxide (DMSO) to obtain a 1 mM stock solution. For proteomics, 4x10^4^ cells were seeded in F-bottom 96-well plates (Greiner bio-one) one day before compound treatment (10^6^ cells per well in 6-well plates for Ubiquitinomics). For proteomics experiments, we used three replicates and four replicates for Ubiquitinomics-related treatment experiments. Cells were treated with either DMSO or the indicated compounds (1 µM) for 30 minutes (Ubiquitinomics) or six hours (proteomics). Following treatment, cells were washed with PBS, and lysed with sodium deoxycholate (SDC) lysis buffer, as reported previously^49^.

### Ubiquitinomics - sample preparation for ubiquitinomics workflows

Ubiquitinomics sample preparation was done according to our recently established protocol with a few modifications^49^. No proteasome inhibitors (such as MG-132) were added to the cells. In brief, protein concentrations were determined using the BCA assay (Merck-Millipore) and proteins digested overnight at 37°C using 100:1 protein:trypsin ratio (Promega). After digestion, immunoprecipitation (IP) buffer (50 mM MOPS pH 7.2, 10 mM Na_2_HPO_4_, 50 mM NaCl) was added to the samples together with K-GG antibody-bead conjugate, followed by a 2 h incubation in a mixer. Bead washing and peptide elution was performed according to manufacturer’s instructions. The peptide eluates were desalted using in-house prepared, 200 µl two-plug C18 StageTips (3M EMPORE)^67^.

For the Ubiquitinomics experiment testing various input amounts, 200 µg of total protein was digested, followed by K-GG peptide enrichment. The K-GG peptide eluate was then sequentially diluted to mimic 50 µg or less protein input per sample. Each sample was measured on LC-MS in triplicate.

### Ubiquitinomics - sample preparation for proteomics upon drug treatment

Cells were washed and lysed in 96-well plates using SDC buffer^49^. After tryptic digestion of proteins (200 ng trypsin/well), the SDC was precipitated using trifluoroacetic acid (TFA) (1% (v/v) final). Peptides were desalted on C18 StageTips^67^. After drying in a vacuum centrifuge (Eppendorf), peptides were resuspended in 0.1% (v/v) TFA and their concentration estimated on a Nanodrop spectrophotometer (Thermo). Eight hundred ng of peptides were loaded per LC-MS acquisition.

### Ubiquitinomics LC-MS Measurement

Peptides were loaded on a 35 cm reversed phase column (75 µm i.d., packed in-house with ReproSil Saphir C18 1.5 µm resin (ReproSil Saphir, Dr. Maisch GmbH)). The column temperature was maintained at 55°C using a column oven. A Vanquish Neo UHPLC system (ThermoFisher) was directly coupled online with the timsTOF HT mass spectrometer (Bruker) via a nano-electrospray source, and peptides were separated with a binary buffer system of buffer A (0.1% FA) and buffer B (80% ACN + 0.1% FA), at a flow rate of 300 nl/min.

For the ubiquitinomics experiment on various input amounts we used either Slice-PASEF or dia-PASEF using a 45-minute linear gradient. For proteomics upon drug-treatment, we used a 60-min gradient while operating the mass spectrometer in dia-PASEF mode. Here, the acquisition method consisted of a MS1 scan, followed by 24x MS2 scans (two ion mobility windows, 100 ms of accumulation/ramp time) with variable window sizes^18^. Ubiquitinomics samples upon drug-treatment were acquired using the 45-minute linear gradient and the 1-Frame Slice-PASEF method only.

### Slice-PASEF module in DIA-NN

To enable Slice-PASEF data analysis, we have incorporated a new module in our DIA-NN software suite. Slice-PASEF data is characterised by several key differences from dia-PASEF, which necessitated dedicated data processing approaches.

First, while dia-PASEF data is processed similarly to regular DIA, in the sense that each precursor ion query is matched to MS/MS spectra acquired with isolation windows containing this query, in Slice-PASEF each frame spans a much broader precursor *m/z* range. Therefore, the distinction of whether or not a precursor needs to be queried against a particular frame is defined by the expected ion mobility value of the precursor, in addition to its *m/z* value. To enable computationally efficient analysis of Slice-PASEF data, we therefore incorporated another layer of information in the internal representation of spectra in DIA-NN, wherein each peak is associated not only with the values of its *m/z*, intensity and ion mobility, but also with the isolation window boundaries, used to obtain it. Thus, for each precursor and frame of interest, DIA-NN first determines if any of the peaks in the fragment spectrum could potentially originate from the precursor ion, given its *m/z* value and predicted ion mobility, and only in case of a possible match compares the fragment ions associated with the precursor ion to the spectrum. In this case, only peaks with matching isolation window boundaries are considered.

Second, in the case of multi-frame Slice-PASEF, DIA-NN needs to consider multiple frames in each subcycle. In this case, DIA-NN separately analyses each fragment ion of a precursor, by comparing it to all the frames of the subcycle and identifying the frame with the most intense matching peak. The intensity of this peak is then regarded as the intensity of the fragment ion in the respective subcycle.

Third, the low number of distinct frames constituting a full DIA cycle in Slice-PASEF also means that it achieves high numbers of data points per peptide peak. Aiming to maintain a minimum of circa three data points per peak at FWHM (full width half maximum) on average, we therefore were able to introduce frame repeats for the 1F and 2F methods, wherein each frame is acquired several times in succession. DIA-NN then merges the spectra corresponding to these repeats prior to performing 2D-peak picking^14^, thus improving the signal-to-noise ratio and increasing sensitivity, eventually inferring the *m/z* and ion mobility of observed ions with better accuracy.

### Raw LC-MS data processing in DIA-NN - General

The data were processed using DIA-NN 1.9.2 (all datasets other than plexDIA) and 2.0.2 (plexDIA data only), downloadable under https://github.com/vdemichev/DiaNN/releases. The majority of settings were kept constant for all data sets processed: Mass accuracies were fixed to 15 ppm (both MS1 and MS2), and the scan windows were set to 6 (analytical flow-rate chromatography), 7 (microliter flow-rate chromatography on the Evosep One System), or 0 (i.e., automatic inference, as per default) for all other datasets. The “--tims-scan” option was supplied to DIA-NN 1.9.2 for the analysis of all Slice-PASEF acquisitions, while 2.0.2 detects these as Slice-PASEF automatically.

### Raw LC-MS data processing in DIA-NN - Benchmark

For the K562 dilution series benchmark dataset, the match-between-runs feature (MBR) was disabled and the previously described empirical DIA-based library was used^20^. Protein inference was disabled to use the protein grouping already present in the spectral library. The spectral library^14^ that was used to analyse the HeLa 0.2 ng and 1 ng acquisitions on the Evosep One system was first refined using an analysis of five 5 ng HeLa acquisitions, with the Library generation strategy set to IDs, RT & IM profiling. The Legacy quantification module was used.

### Raw LC-MS data processing in DIA-NN - FACS SCP

For U2OS single-cell proteomics experiments, a fully predicted spectral library was generated using the Uniprot Proteome UP000005640 as input (i.e., without isoforms; retrieved 14.01.2025) while allowing for one missed cleavage, a precursor *m/z* range from 300-1800 *m/z* at precursor charge states 2 and 3. Common contaminants were added as defined by DIA-NN. All Cysteine residues were set as carbamidomethylated; no variable modifications were allowed. The LC-MS data was searched using this library using mass accuracies set to 15 ppm (both MS1 and MS2) and else under default settings, with protein inference set to gene-level and the MBR feature enabled. The quantification used the Legacy (direct) strategy.

### Raw LC-MS data processing in DIA-NN - plexDIA SCP

For plexDIA data analysis, we first added the mTRAQ d0-channel to the predicted spectral library from above (”U2OS single-cell proteomics experiments”) using DIA-NN’s deep-learning peptide prediction module and the custom commands: --fixed-mod mTRAQ,140.0949630177,nK / --lib-fixed-mod mTRAQ / --original-mods / --report-lib-info. The obtained library was subsequently refined using the three 100-cell-equivalent (see above) plexDIA acquisitions with 15 ppm MS1 and MS2 mass accuracy, no MBR, inference set to the gene-level and the custom commands: --fixed-mod mTRAQ,140.0949630177,nK / --original-mods / --report-lib-info. Third, the single-cell analyses or two bulk measurements were searched using the same settings but MBR enabled and inference turned off since unnecessary at this point. Custom commands were: --fixed-mod mTRAQ,140.0949630177,nK / --channels mTRAQ,0,nK,0:0; / mTRAQ,4,nK,4.0070994:4.0070994;mTRAQ,8,nK,8.0141988132:8.0141988132 / --peak-translation / --original-mods / --report-lib-info / --tims-scan / --channel-spec-norm. Note that “-- channel-spec-norm” is essential for this type of data to benefit from channel-specific Q.Value assessment and normalization.The MBR feature was enabled to increase comparability between samples despite using the 100-cell equivalent empirical library, as this resulted in higher data completeness. Quantification employed the QuantUMS algorithm in high-precision mode^68^.

### Raw LC-MS data processing in DIA-NN - Ubiquitinomics

For the low-input titration Ubiquitinomics experiment, we generated a fully predicted spectral library using the Uniprot reference proteome (UP00005640), including one missed cleavage, two variable modifications allowing for Methionine oxidation, protein N-terminal acetylation and K-GG-remnants on Lysine residues. Precursor charge-states of 2 and 3 were allowed while all other options were kept default. For all searches, mass accuracies were fixed to 15 ppm (both MS1 and MS2) while the scan window was inferred automatically. This library was refined using four dia-PASEF LC-MS acquisitions of a HEK cell line digest on the same LC-MS setup including MBR and peptidoform scoring in DIA-NN. Using this library on our full titration series data, we used the Legacy quantification mode with MBR disabled and gene-level inference.

For the compound-treatment Proteomics experiments data, we used a predicted spectral library similar to the defined above (Ubiquitinomics dataset) but without allowing variable modifications. Gene-level inference and MBR was enabled and quantification leveraged QuantUMS in high-precision mode^68^.

The data from compound-treatment Ubiquitinomics experiments was searched using the refined library from the titration experiment with peptidoform scoring, inference on gene-level, and the high-precision setting for quantification by QuantUMS^68^.

### Data Analysis - General

Data post-processing and analysis used Python 3.9 in conjunction with the pandas 2.2.3, numpy 2.1.3, and scipy 1.14.1 packages, as well as R 4.3.1 with the dplyr package. DIA-NN reports were filtered for 1% FDR on precursor, protein or gene level.

Coefficients of variation (CVs) were obtained by dividing the standard deviation of non-log-transformed signal intensities with n - 1 degrees of freedom by their mean and only reported for identifications with n >= 3 if not indicated otherwise. CVs are given as values or percentages as indicated. All plots were generated in Python using the Seaborn 0.13.2 package and arranged using Inkscape 1.2 and 1.4.

### Data Analysis - Benchmark Dataset

For benchmarks, FDR was applied per LC-MS run. When using a spectral library from DIA-data, data was filtered to the corresponding Library q-values.

### Data Analysis - Single-cell Proteomics FACS Dataset

To compare MS1 and MS2 precursor intensities from FACS-sorted cells, we first extracted the mean normalised and log_2_-converted quantities from DIA-NN for each protein group (i..e, “Ms1.Area” and “Precursor.Quantity”) and method. Following the subtraction of their individual medians for both, MS1 and MS2, we plotted these intensities in a contour plot based on a gaussian kernel density estimation using the seaborn kdeplot function with five levels and else under default settings.

### Data Analysis - Single-cell Proteomics plexDIA Dataset

The data report from plexDIA experiment data was filtered to less or equal than 1% FDR on Lib.PG.Q.Value and Global.PG.Q.Value, as well as to less or equal than 5% on PG.Q.Value and Channel.Q.Value. Two LC-MS acquisitions used the bulk samples and 76 runs contained single cells pooled as 3-plex or 2-plex summing to a total of single 210 cells (109x Jurkat, 101x U937) plus nine empty control channels. Upon filtering for failed cell preparations and removing outlier channels flagged by the elevated presence of contaminants we arrived at a total of 191 single cells (101x Jurkat, 90x U937) from 75 LC-MS runs. Log_2_-transformed normalised precursor quantities (“Precursor.Normalised”) were first median centered per single-cell-channel considering only those precursors with 100% completeness and then summarised to MaxLFQ protein quantities using the iq R package^69^. The protein quantity matrix filtered to 100% data completeness was submitted to PCA analysis using R’s svd() function.

To extract biological processes differing between cell types, we applied gene set enrichment analysis (GSEA). Here, normalised cell-type specific gene quantities (see above) were filtered to 10% overall completeness, leading to 1516 entries. To score gene products for a pre-ranked GSEA, we calculated a ranking metric based on the Mann–Whitney U p-value and the direction of change:

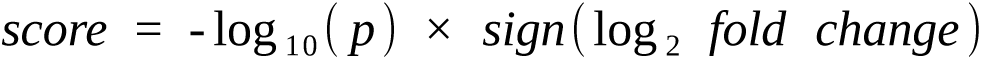

Functional enrichment was then performed using the ranked gene list via the GSEApy package (with the “KEGG human 2021” reference gene set while employing 1000 multiple testing correction permutations, and setting sizes of individual gene sets to contain between 8 and 500 members^70,71^. The output was filtered allowing a maximum false-discovery rate q-value of 50%.

For correlating histone protein abundances with U937 cell sizes, histone protein quantities obtained in >70% of measurements were averaged upon minimum value imputation per protein group. The data was then split to three equally sized bins based on normalised NUCKS1-abundance. We performed Student’s t-test followed by multiple testing correction via the Benjamini-Hochberg procedure between the high and low tertile groups to identify proteins significantly changing abundance levels along with NUCKS1 using a 10% FDR cut- off. Proteins were assigned to cell-cycle state(s) based on proteomics data from Altun and coworkers^41^.

### Data Analysis - Ubiquitinomics Dataset

Ubiquitinomics data was filtered to a Lib.PG.Q.Value of less or equal than 1%. K-GG-remnant peptide-centric analyses used the Modified.Sequence column that was filtered to only contain ubiquitinylated peptides (“UniMod: 121”). Data from compound-treatment experiments was filtered to less or equal than 1% on Lib.PG.Q.Value and PG.Q.Value Level. Quantities for Volcano plots were log_2_-transformed, filtered to 100% (gene-centric Proteomics and Ubiquitinomics) or 75% completeness (K-GG-remnant peptide-centric Ubiquitinomics) per treatment and their statistical significance tested using a two-sample independent Student’s t-test followed by multiple testing correction according to Benjamini-Hochberg^72^. UbiBrowser reference tables for Deubiquitinases (predicted and literature curated lists, downloaded 7.2.2025) were filtered for “Homo sapiens” and Ubiquitin carboxyl-terminal hydrolase 7 (USP7, ‘Q93009’) for comparisons involving FT671 treatments. Ubiquitination sites for the new protein hit MAGED4 (‘Q96JG8’) were manually validated using the linked repositories of PTMeXchange with PRIDE and PeptideAtlas to the UniProt database (accessed, 30.7.2025)^73^.

## Acknowledgements

We acknowledge the Charité Core Facility High Throughput Mass Spectrometry, led by Michael Mülleder, for supporting the K562 dilution series benchmark. This work is funded by the German Ministry of Education and Research (BMBF), as part of the National Research Node “Mass spectrometry in Systems Medicine” (MSCoreSys), under grant agreement 161L0221 (to V.D.) and 031L0220 (to M.R.) and 161L0222 (F.C. and D.Q.). Further support was given from the European Research Council (ERC-SyG-2020951475) (to M.R.), a Bits to Bytes award from MLSC (to N.S.), and a MIRA award from the NIGMS of the NIH (R35GM148218) (to N.S.).

## Author contributions

V.D., L.S. and N.N. conceived the study. L.K. cultivated Jukat and U937 cells. N.S. designed the plexDIA experiment. L.S., K.L., K.F., F.M., L.K., A.L., M.S., D.Q., and L.R.S. prepared samples and collected and processed raw data. V.D. developed the raw data processing algorithms. L.R.S. and J. G. performed data analysis and generated plots. L.R.S. and V.D. wrote the drafts; all authors contributed to writing the manuscript. V.D., M.S., N.S., M.R. and F.C. supervised the study.

## Data and Code Availability

Slice-PASEF and dia-PASEF mass spectrometry data, DIA-NN logs, pipelines and reports have been deposited via the PRIDE partner repository ProteomeXchange^74^ and can be accessed using the dataset identifiers PXD063201 (Benchmark), PXD063234 (FACS U2OS), PXD063280 (plexDIA), and PXD063371 (Ubiquitinomics).

The interactive Shiny application to generate Slice-PASEF methods is available on Zenodo (DOI: 10.5281/zenodo.17034921).

---

**Fig. S1:**
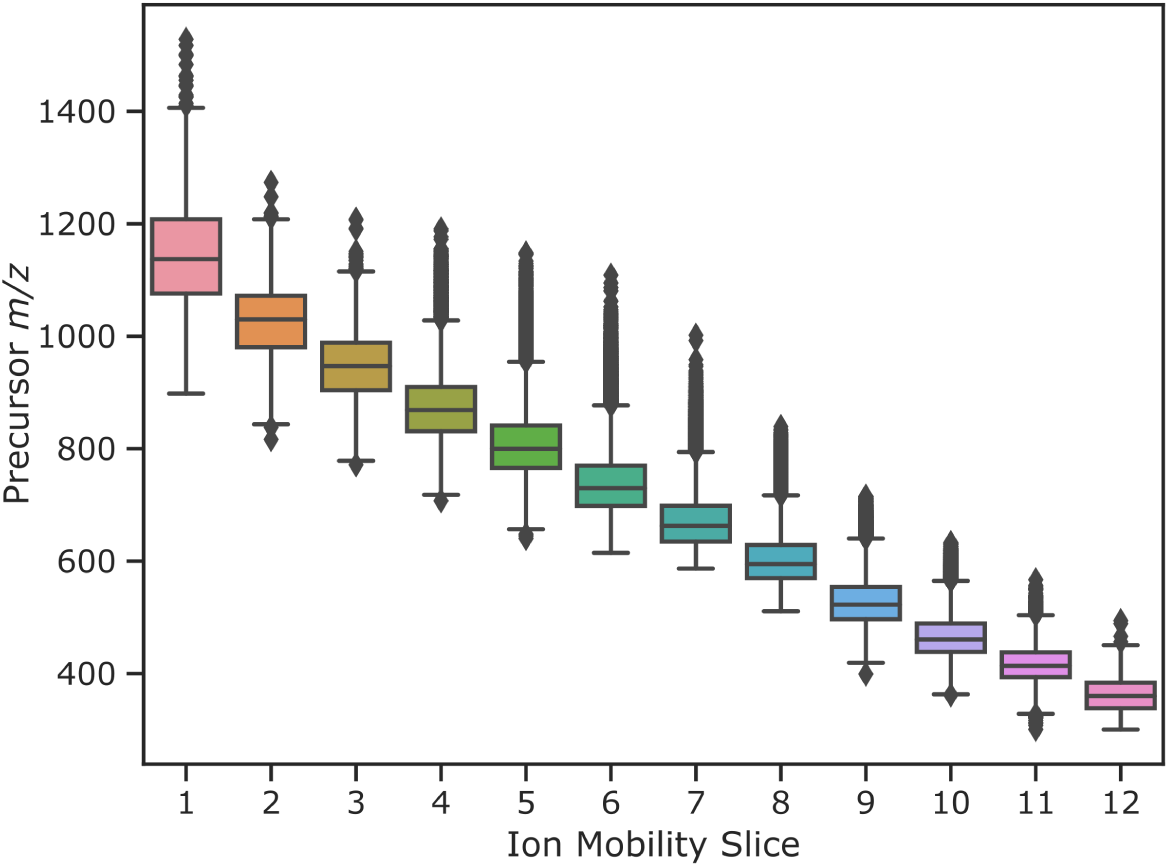
Distribution of precursor m/z along ion mobility slices Boxplots on precursor ion distributions in dia-PASEF data from Meier and coworkers13. Data was filtered to the ion mobility range of interest (m/z 350-1050 & 0.75-1.15 V*s/cm2) and split into twelve bins using 0.0375 V*s/cm2 intervals. The interquartile range (IQR) (i.e., 25-75% of data) equals 71.18 m/z (±24.48 s.d.) and decreases along the slices (towards lower ion mobility). Whiskers span over 1.5 x IQR. Outliers are shown as black diamonds.

**Fig. S2:**
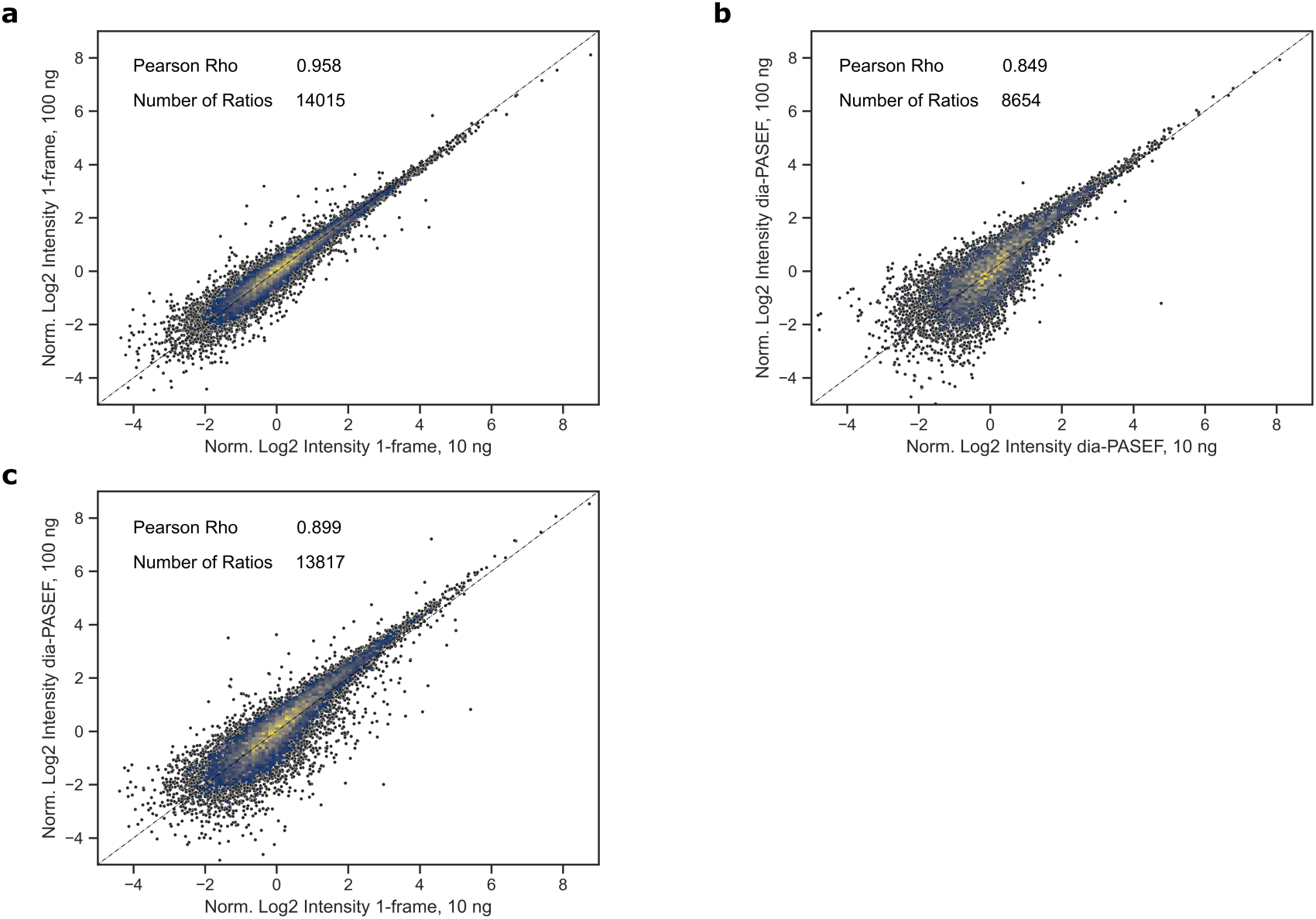
Quantitative similarity between dia-PASEF and 1-Frame Slice-PASEF methods on analytical flow rate LC-MS Normalised log_2_-transformed signal intensities of shared precursor identifications are shown for different method combinations and injection amounts. a) Comparison of 10 and 100 ng K562 loads using 1F Slice-PASEF data. b) Comparison of 10 and 100 ng K562 loads using dia-PASEF data. c). Comparison of 100 ng dia-PASEF vs. 10 ng 1F Slice-PASEF data, both processed in independent DIA-NN searches. Input quantities represent the median from three technical replicates.

**Fig. S3:**
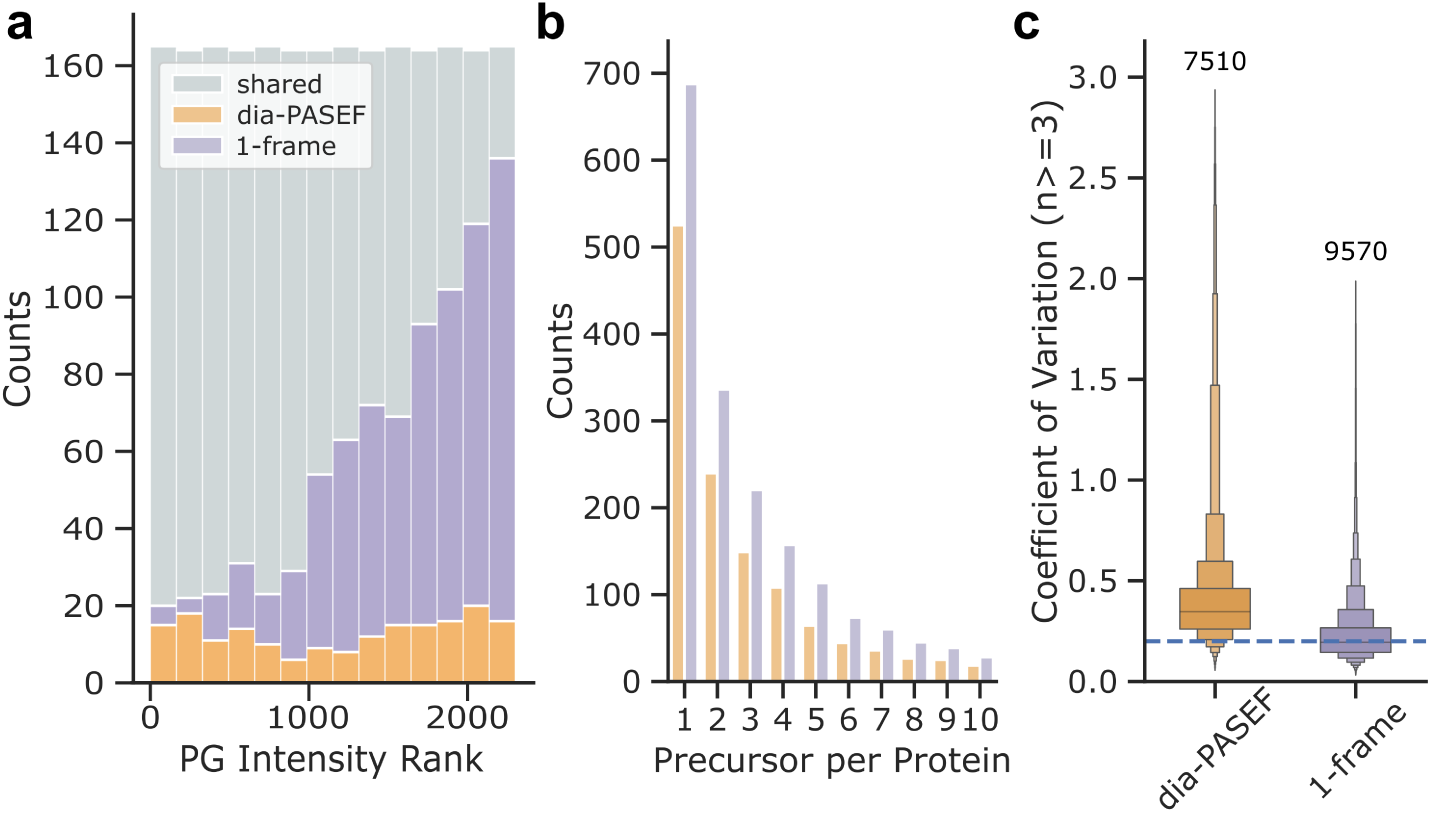
Figures of merit for dia-PASEF and 1-Frame Slice-PASEF methods comparison on U2OS single-cell proteomics a) Proteome coverage by intensity-binned protein groups from both methods. Identifications of dia- PASEF are shown in orange, 1-Frame Slice-PASEF in purple, and shared identifications in grey. b) Average identified precursors per protein for both methods. c) Quantitative precision for identified precursors of both methods as Boxenplot. Medians (dia-PASEF: 0.347 / 1-Frame Slice-PASEF: 0.195) and 50% IQRs are highlighted by the central box while each quantile further outwards represents half of the remaining fraction. Outliers (i.e. upper and lower 0.7%) are not shown. The sample size for each method is given above the boxes. The dashed line marks a CV of 20%.

**Fig. S4:**
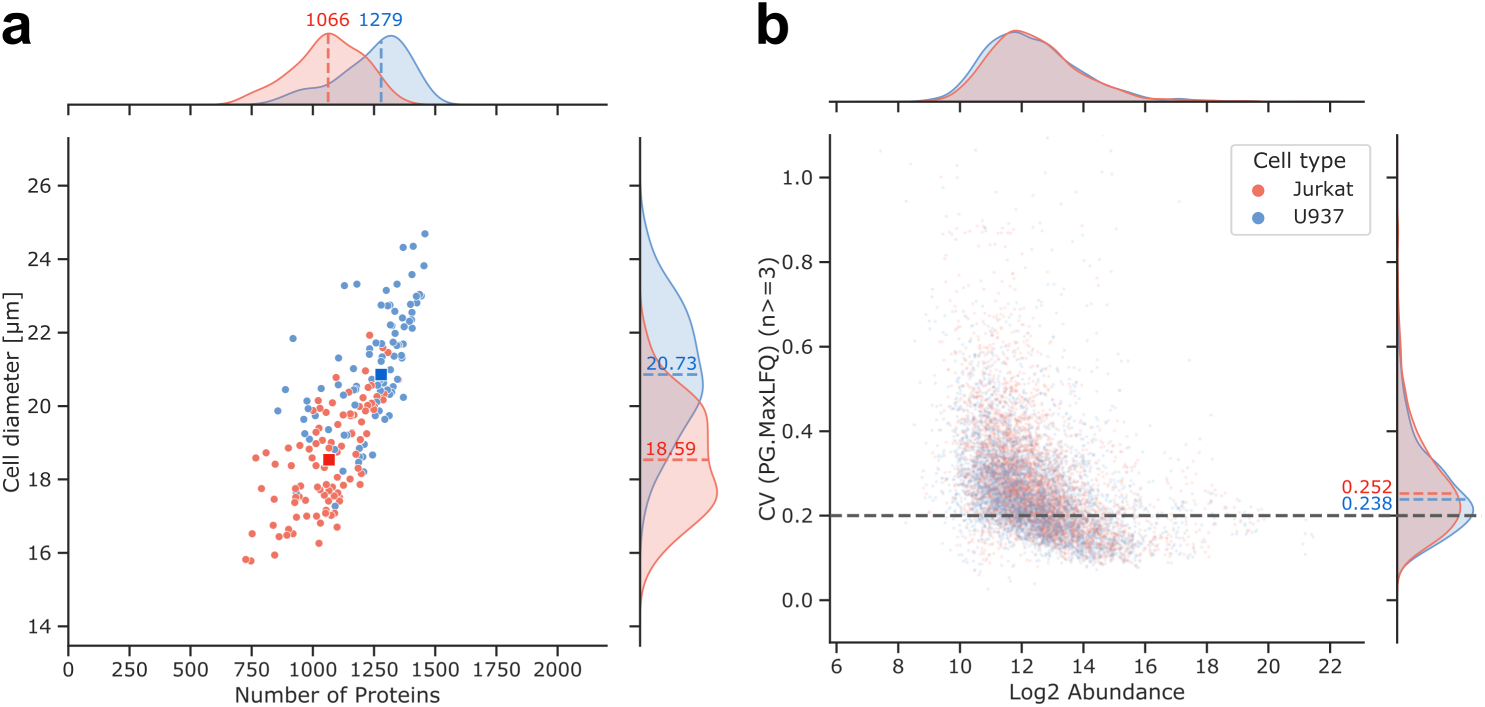
Overall identifications in plexDIA Slice-PASEF and their relationship to cell size and quantitative precision (a) The number of identified protein groups increases with cell size. Medians per cell type are highlighted by a darker rectangle for each cell type with the corresponding values and distributions being indicated at the axis margins. b) Quantitative precision measured by the coefficient of variation (CV) per protein group across the quantitative domain. The dashed black line marks a CV of 20% while the dashed lines at the right axis margin mark the medians for each cell type (Jurkat: 25.2% / U937: 23.8%). The colour code for each cell type is indicated in panel (b).

**Fig. S5:**
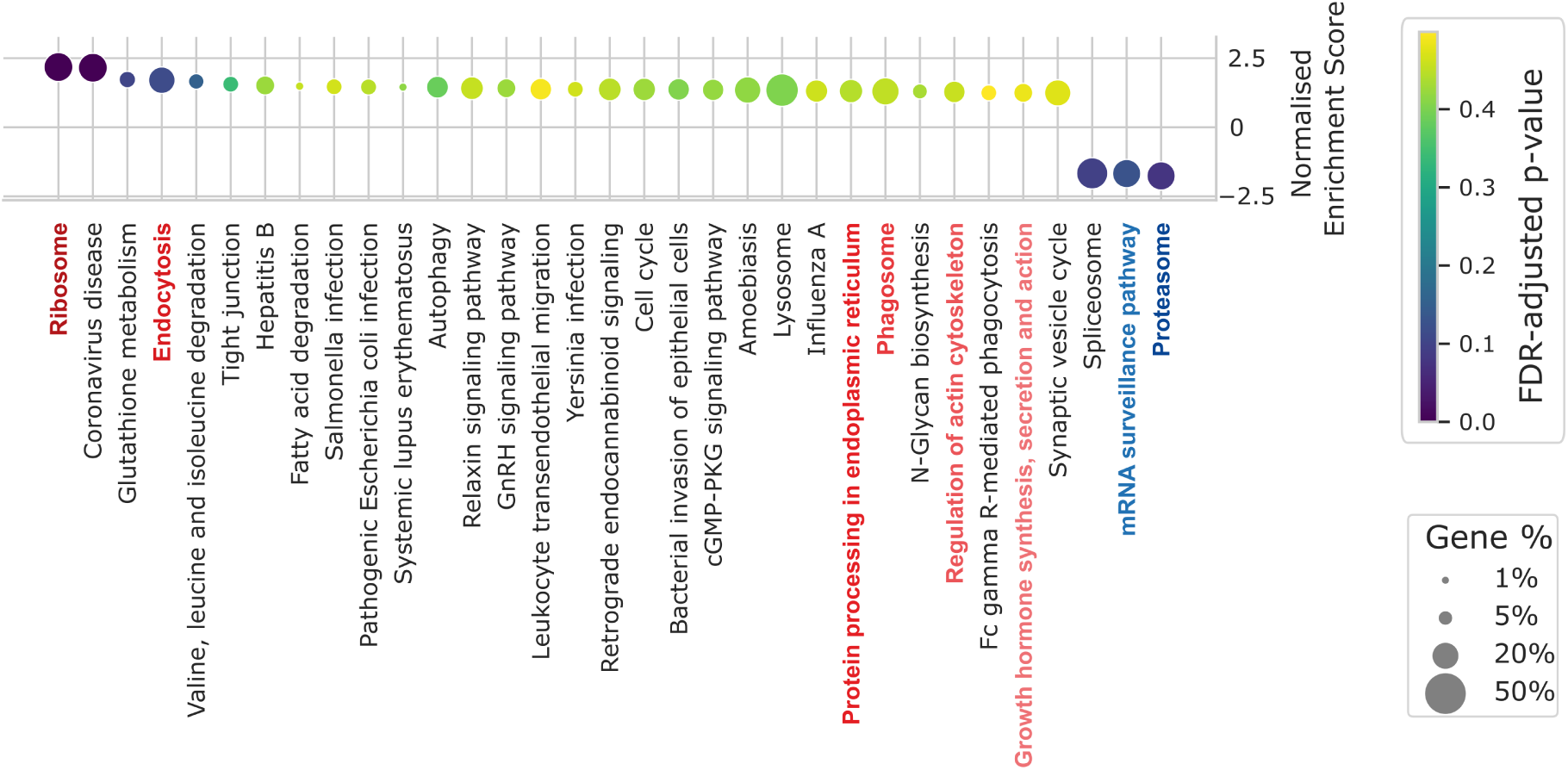
Differential protein abundances in biological processes characteristic to Jurkat and U937 cell lines Gene set enrichment analysis (GSEA) on quantitatively different shared gene products of both cell types. Upon applying a 10% completeness filter, gene products were ranked by a score derived from Mann-Whitney-U p-values and the direction of fold-change (Methods) before forwarding to pre-ranked GSEA. Terms passing a 50% FDR q-value threshold are displayed with declining normalised enrichment scores. The colour hue illustrates multiple testing-adjusted p-values while the size of each sign symbolizes the proportion of genes at the enrichment score peak as defined by the GSEApy Python package. Selected gene sets are highlighted in red for U937 and in blue for Jurkat cells.

**Fig. S6:**
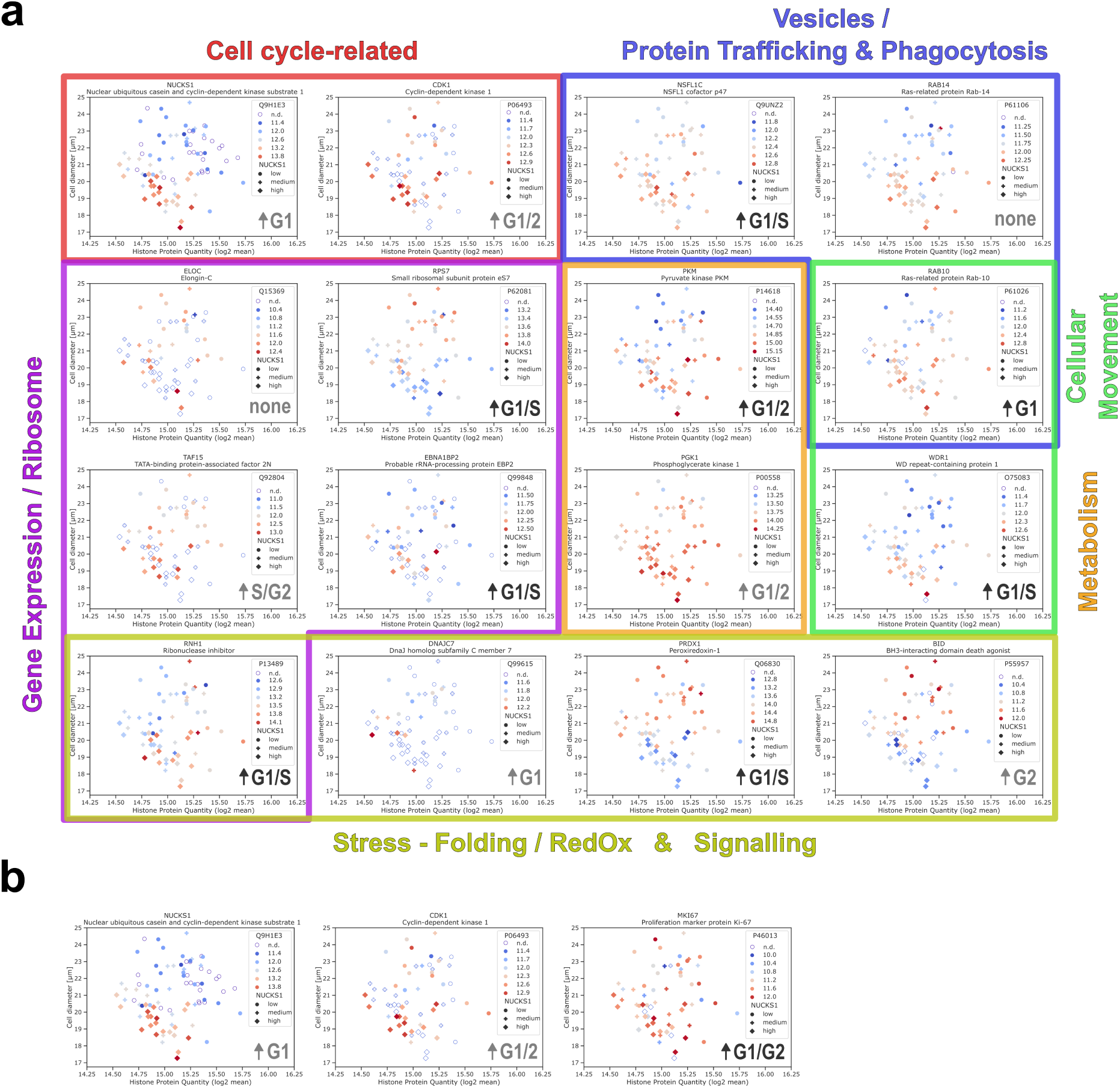
Quantitative protein abundance differences between NUCKS1-stratified groups in U937 cells The log2-converted, normalised and averaged quantities of histone proteins are compared with observed cell sizes, separated into abundance tertiles of the NUCKS1 gene product (not detected, n.d.: hollow circle, low: circle, medium: cross, high: diamond). The colour hue illustrates the corresponding quantities of a protein of interest, as indicated at the top right of each subpanel. Panel (a) shows the significant hits (10% FDR cut-off, from performing a Student’s t-test and Benjamini-Hochberg multiple testing correction) from comparing the proteins between the low and high NUCKS1 abundance tertiles. Proteins were manually associated with broad biological categories, as indicated by border and label in the same colour. Suggested cell-cycle stages associated with higher levels of each gene product were inferred from data by Altun and colleagues41 and are highlighted in black if ≤10% FDR therein. Panel (b) compares NUCKS1 with two cell-cycle marker proteins, CDK1 (also present in panel a) and MKI67, previously highlighted as cell cycle markers by Bubis and colleagues39. Data were processed as for panel (a).

**Fig. S7:**
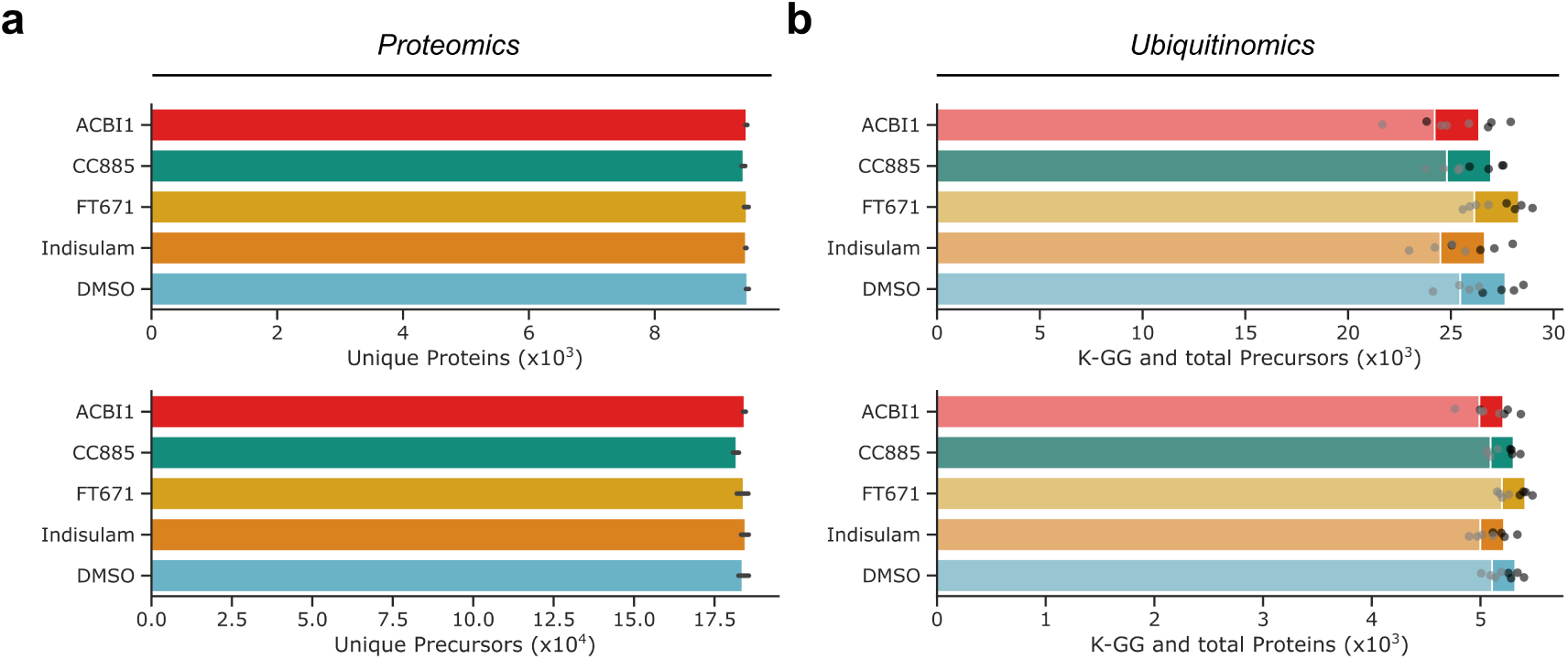
Proteomics and Ubiquitinomics Identifications upon drug treatment (a) Protein group and precursor identification numbers in drug treatment experiment using dia-PASEF proteomics. Error bars span the 95% confidence interval. b) K-GG remnant peptide precursors and ubiquitylated protein group identification numbers in Ubiquitinomics drug treatment experiment using 1- Frame Slice-PASEF. Lighter colours symbolize K-GG-remnant entities while darker colours indicate unmodified precursor or protein group identifications. Individual values are shown as jittered points. DMSO represents the mock treatment.

## References

1. Zhang, F., Ge, W., Ruan, G., Cai, X. & Guo, T. Data-independent acquisition mass spectrometry-based proteomics and software tools: A glimpse in 2020. Proteomics 20, e1900276 (2020).

2. Fröhlich, K. et al. Data-independent acquisition: A milestone and prospect in clinical mass spectrometry-based proteomics. Mol. Cell. Proteomics 23, 100800 (2024).

3. Messner, C. B. et al. Ultra-High-Throughput Clinical Proteomics Reveals Classifiers of COVID-19 Infection. Cell Syst 11, 11–24.e4 (2020).

4. Muenzner, J. et al. Natural proteome diversity links aneuploidy tolerance to protein turnover. Nature 630, 149–157 (2024).

5. Mi, Y. et al. High-throughput mass spectrometry maps the sepsis plasma proteome and differences in patient response. Sci. Transl. Med. 16, eadh0185 (2024).

6. Brunner, A.-D. et al. Ultra-high sensitivity mass spectrometry quantifies single-cell proteome changes upon perturbation. Mol. Syst. Biol. 18, e10798 (2022).

7. Mund, A. et al. Deep Visual Proteomics defines single-cell identity and heterogeneity. Nat. Biotechnol. 40, 1231–1240 (2022).

8. Wei, L., Meyer, J. G. & Schilling, B. Quantification of site-specific protein lysine acetylation and succinylation stoichiometry using data-independent acquisition mass spectrometry. J. Vis. Exp. e57209 (2018) doi:10.3791/57209.

9. Rosenberger, F. A., Thielert, M. & Mann, M. Making single-cell proteomics biologically relevant. Nat. Methods 20, 320–323 (2023).

10. Ghosh, G., Shannon, A. E. & Searle, B. C. Data acquisition approaches for single cell proteomics. Proteomics 25, e2400022 (2025).

11. Gatto, L. et al. Initial recommendations for performing, benchmarking and reporting single-cell proteomics experiments. Nat. Methods 20, 375–386 (2023).

12. Slavov, N. Driving Single Cell Proteomics Forward with Innovation. J. Proteome Res. 20, 4915–4918 (2021).

13. Meier, F. et al. diaPASEF: parallel accumulation–serial fragmentation combined with data-independent acquisition. Nat. Methods 17, 1229–1236 (2020).

14. Demichev, V. et al. dia-PASEF data analysis using FragPipe and DIA-NN for deep proteomics of low sample amounts. Nat. Commun. 13, 3944 (2022).

15. Distler, U. et al. midiaPASEF maximizes information content in data-independent acquisition proteomics. bioRxiv 2023.01.30.526204 (2023) doi:10.1101/2023.01.30.526204.

16. Skowronek, P. et al. Synchro-PASEF allows precursor-specific fragment ion extraction and interference removal in data-independent acquisition. Mol. Cell. Proteomics 22, 100489 (2023).

17. Mendes, M. L., Borrmann, K. F. & Dittmar, G. Eleven shades of PASEF. Expert Rev. Proteomics (2024) doi:10.1080/14789450.2024.2413092.

18. Skowronek, P. et al. Rapid and in-depth coverage of the (phospho-)proteome with deep libraries and optimal window design for dia-PASEF. Mol. Cell. Proteomics 21, 100279 (2022).

19. Derks, J. et al. Increasing the throughput of sensitive proteomics by plexDIA. Nat. Biotechnol. (2022) doi:10.1038/s41587-022-01389-w.

20. Szyrwiel, L., Gille, C., Mülleder, M., Demichev, V. & Ralser, M. Fast proteomics with dia-PASEF and analytical flow-rate chromatography. Proteomics 24, e2300100 (2024).

21. Derks, J., et al. Single-nucleus proteomics identifies regulators of protein transport. bioRxiv (2024) doi:10.1101/2024.06.17.599449.

22. Meier, F. et al. Parallel Accumulation–Serial Fragmentation (PASEF): Multiplying Sequencing Speed and Sensitivity by Synchronized Scans in a Trapped Ion Mobility Device. J. Proteome Res. 14, 5378–5387 (2015).

23. Geiger, T., Cox, J. & Mann, M. Proteomics on an orbitrap benchtop mass spectrometer using all-ion fragmentation. Mol. Cell. Proteomics 9, 2252–2261 (2010).

24. Silveira, J. A., Ridgeway, M. E., Laukien, F. H., Mann, M. & Park, M. A. Parallel accumulation for 100% duty cycle trapped ion mobility-mass spectrometry. Int. J. Mass Spectrom. 413, 168–175 (2017).

25. Amodei, D. et al. Improving precursor selectivity in data-independent acquisition using overlapping windows. J. Am. Soc. Mass Spectrom. 30, 669–684 (2019).

26. Willems, S., Voytik, E., Skowronek, P., Strauss, M. T. & Mann, M. AlphaTims: Indexing trapped ion mobility spectrometry-TOF data for fast and easy accession and visualization. Mol. Cell. Proteomics 20, 100149 (2021).

27. Kulak, N. A., Pichler, G., Paron, I., Nagaraj, N. & Mann, M. Minimal, encapsulated proteomic-sample processing applied to copy-number estimation in eukaryotic cells. Nat. Methods 11, 319–324 (2014).

28. Derks, J. & Slavov, N. Strategies for Increasing the Depth and Throughput of Protein Analysis by plexDIA. J. Proteome Res. 22, 697–705 (2023).

29. Leduc, A., Khoury, L., Cantlon, J., Khan, S. & Slavov, N. Massively parallel sample preparation for multiplexed single-cell proteomics using nPOP. Nat. Protoc. 19, 3750– 3776 (2024).

30. Barber, E. K., Dasgupta, J. D., Schlossman, S. F., Trevillyan, J. M. & Rudd, C. E. The CD4 and CD8 antigens are coupled to a protein-tyrosine kinase (p56lck) that phosphorylates the CD3 complex. Proc. Natl. Acad. Sci. U. S. A. 86, 3277–3281 (1989).

31. Dietrich, J., Hou, X., Wegener, A. M. & Geisler, C. CD3 gamma contains a phosphoserine-dependent di-leucine motif involved in down-regulation of the T cell receptor. EMBO J. 13, 2156–2166 (1994).

32. Horwood, N. J. et al. Bruton’s tyrosine kinase is required for TLR2 and TLR4-induced TNF, but not IL-6, production. J. Immunol. 176, 3635–3641 (2006).

33. Mohamed, A. J. et al. Bruton’s tyrosine kinase (Btk): function, regulation, and transformation with special emphasis on the PH domain. Immunol. Rev. 228, 58–73 (2009).

34. Lehmann, M. H. Recombinant human granulocyte-macrophage colony-stimulating factor triggers interleukin-10 expression in the monocytic cell line U937. Mol. Immunol. 35, 479–485 (1998).

35. Chanput, W., Peters, V. & Wichers, H. THP-1 and U937 Cells. in The Impact of Food Bioactives on Health 147–159 (Springer International Publishing, Cham, 2015). doi:10.1007/978-3-319-16104-4_14.

36. Ternette, N. et al. Early kinetics of the HLA class I-associated peptidome of MVA.HIVconsv-infected cells. J. Virol. 89, 5760–5771 (2015).

37. Orsburn, B. C. Single cell proteomics by mass spectrometry reveals deep epigenetic insight into the actions of an orphan histone deacetylase inhibitor. bioRxiv 2024.01.05.574437 (2024) doi:10.1101/2024.01.05.574437.

38. Hume, S. et al. The NUCKS1-SKP2-p21/p27 axis controls S phase entry. Nat. Commun. 12, 6959 (2021).

39. Bubis, J. A. et al. Challenging the Astral mass analyzer to quantify up to 5,300 proteins per single cell at unseen accuracy to uncover cellular heterogeneity. Nat. Methods 1–10 (2025) doi:10.1038/s41592-024-02559-1.

40. Lanz, M. C., Fuentes Valenzuela, L., Elias, J. E. & Skotheim, J. M. Cell size contributes to single-cell proteome variation. J. Proteome Res. 22, 3773–3779 (2023).

41. Herr, P. et al. Cell Cycle Profiling Reveals Protein Oscillation, Phosphorylation, and Localization Dynamics*. Mol. Cell. Proteomics 19, 608–623 (2020).

42. Hammarén, H. M., Geissen, E.-M., Potel, C. M., Beck, M. & Savitski, M. M. Protein-Peptide Turnover Profiling reveals the order of PTM addition and removal during protein maturation. Nat. Commun. 13, 7431 (2022).

43. Leijten, N. M., Heck, A. J. R. & Lemeer, S. Histidine phosphorylation in human cells; a needle or phantom in the haystack? Nat. Methods 19, 827–828 (2022).

44. Kaulich, P. T., Jeong, K., Kohlbacher, O. & Tholey, A. Influence of different sample preparation approaches on proteoform identification by top-down proteomics. Nat. Methods 21, 2397–2407 (2024).

45. Prus, G., Satpathy, S., Weinert, B. T., Narita, T. & Choudhary, C. Global, site-resolved analysis of ubiquitylation occupancy and turnover rate reveals systems properties. Cell (2024) doi:10.1016/j.cell.2024.03.024.

46. Rankovic, Z. et al. Unbiased mapping of cereblon neosubstrate landscape by high-throughput proteomics. bioRxiv 2024.10.18.618633 (2024) doi:10.1101/2024.10.18.618633.

47. Tsai, J. M., Nowak, R. P., Ebert, B. L. & Fischer, E. S. Targeted protein degradation: from mechanisms to clinic. Nat. Rev. Mol. Cell Biol. 25, 740–757 (2024).

48. Hansen, F. M. et al. Data-independent acquisition method for ubiquitinome analysis reveals regulation of circadian biology. Nat. Commun. 12, 254 (2021).

49. Steger, M. et al. Time-resolved in vivo ubiquitinome profiling by DIA-MS reveals USP7 targets on a proteome-wide scale. Nat. Commun. 12, 5399 (2021).

50. Farnaby, W. et al. BAF complex vulnerabilities in cancer demonstrated via structure-based PROTAC design. Nat. Chem. Biol. 15, 672–680 (2019).

51. Han, T. et al. Anticancer sulfonamides target splicing by inducing RBM39 degradation via recruitment to DCAF15. Science 356, (2017).

52. Matyskiela, M. E. et al. A novel cereblon modulator recruits GSPT1 to the CRL4(CRBN) ubiquitin ligase. Nature 535, 252–257 (2016).

53. Wang, X. et al. UbiBrowser 2.0: a comprehensive resource for proteome-wide known and predicted ubiquitin ligase/deubiquitinase-substrate interactions in eukaryotic species. Nucleic Acids Res. 50, D719–D728 (2022).

54. Ye, Z. et al. Enhanced sensitivity and scalability with a Chip-Tip workflow enables deep single-cell proteomics. Nat. Methods 1–11 (2025) doi:10.1038/s41592-024-02558-2.

55. Steger, M., Karayel, Ö. & Demichev, V. Ubiquitinomics: History, methods, and applications in basic research and drug discovery. Proteomics 22, e2200074 (2022).

56. Szyrwiel, L., Sinn, L., Ralser, M. & Demichev, V. Slice-PASEF: fragmenting all ions for maximum sensitivity in proteomics. bioRxiv 2022.10.31.514544 (2022) doi:10.1101/2022.10.31.514544.

57. Skowronek, P., Wallmann, G., Wahle, M., Willems, S. & Mann, M. An accessible workflow for high-sensitivity proteomics using parallel accumulation–serial fragmentation (PASEF). Nat. Protoc. 1–30 (2025) doi:10.1038/s41596-024-01104-w.

58. Khan, S. et al. Inferring post-transcriptional regulation within and across cell types in human testis. bioRxivorg 2024.10.08.617313 (2024) doi:10.1101/2024.10.08.617313.

59. Leduc, A., Harens, H. & Slavov, N. Modeling and interpretation of single-cell proteogenomic data. ArXiv (2023) doi:10.48550/arXiv.2308.07465.

60. Lenz, S. et al. Reliable identification of protein-protein interactions by crosslinking mass spectrometry. Nat. Commun. 12, 1–11 (2021).

61. Wozniak, J. M. et al. Enhanced mapping of small-molecule binding sites in cells. Nat. Chem. Biol. 20, 823–834 (2024).

62. Gómez-Varela, D. et al. Increasing taxonomic and functional characterization of host-microbiome interactions by DIA-PASEF metaproteomics. Front. Microbiol. 14, 1258703 (2023).

63. Oliinyk, D. et al. DiaPASEF analysis for HLA-I peptides enables quantification of common cancer neoantigens. Mol. Cell. Proteomics 24, 100938 (2025).

64. Specht, H., et al. Automated sample preparation for high-throughput single-cell proteomics. bioRxiv 399774 (2018) doi:10.1101/399774.

65. Petelski, A. A. et al. Multiplexed single-cell proteomics using SCoPE2. Nat. Protoc. 16, 5398–5425 (2021).

66. Leduc, A., Huffman, R. G., Cantlon, J., Khan, S. & Slavov, N. Exploring functional protein covariation across single cells using nPOP. Genome Biol. 23, 261 (2022).

67. Rappsilber, J., Mann, M. & Ishihama, Y. Protocol for micro-purification, enrichment, pre-fractionation and storage of peptides for proteomics using StageTips. Nat. Protoc. 2, 1896–1906 (2007).

68. Kistner, F., Grossmann, J. L., Sinn, L. R. & Demichev, V. QuantUMS: uncertainty minimisation enables confident quantification in proteomics. *bioRxiv* 2023.06.20.545604 (2023) doi:10.1101/2023.06.20.545604.

69. Pham, T. V., Henneman, A. A. & Jimenez, C. R. iq: an R package to estimate relative protein abundances from ion quantification in DIA-MS-based proteomics. Bioinformatics 36, 2611–2613 (2020).

70. Kuleshov, M. V. et al. Enrichr: a comprehensive gene set enrichment analysis web server 2016 update. Nucleic Acids Res. 44, W90–7 (2016).

71. Fang, Z., Liu, X. & Peltz, G. GSEApy: a comprehensive package for performing gene set enrichment analysis in Python. Bioinformatics 39, btac757 (2023).

72. Benjamini, Y. & Hochberg, Y. Controlling the false discovery rate: A practical and powerful approach to multiple testing. J. R. Stat. Soc. Series B Stat. Methodol. 57, 289– 300 (1995).

73. UniProt Consortium. UniProt: The universal protein knowledgebase in 2025. Nucleic Acids Res. 53, D609–D617 (2025).

74. Perez-Riverol, Y. et al. The PRIDE database and related tools and resources in 2019: improving support for quantification data. Nucleic Acids Res. 47, D442–D450 (2019).

